# Polymerase Theta Inhibition Kills Homologous Recombination Deficient Tumors

**DOI:** 10.1101/2020.05.23.111658

**Authors:** Jia Zhou, Camille Gelot, Constantia Pantelidou, Adam Li, Hatice Yücel, Rachel E. Davis, Anniina Farkkila, Bose Kochupurakkal, Aleem Syed, Geoffrey I. Shapiro, John A. Tainer, Brian S. J. Blagg, Raphael Ceccaldi, Alan D. D’Andrea

**Affiliations:** Department of Radiation Oncology, Dana-Farber Cancer Institute, Harvard Medical School, Boston, MA 02215, USA; Inserm U830, PSL Research University, Institut Curie, 75005, Paris, France; Department of Medical Oncology, Dana-Farber Cancer Institute and Department of Medicine, Harvard Medical School, Boston, Massachusetts, USA; Department of Chemistry and Biochemistry, University of Notre Dame, Notre Dame, IN 46556, USA; Departments of Cancer Biology and of Molecular and Cellular Oncology, University of Texas MD Anderson Cancer Center, Houston, TX 77030, USA; Center for DNA Damage and Repair, Dana-Farber Cancer Institute, Boston, MA, USA

**Keywords:** Novobiocin, Polymerase theta (POLθ), Homologous Recombination, PARP inhibitor resistance

## Abstract

PARP inhibitors (PARPi) have become a new line of therapy for Homologous Recombination (HR)-deficient cancers. However, resistance to PARPi has emerged as a major clinical problem. DNA polymerase theta (POLθ) is synthetic lethal with HR and a druggable target in HR-deficient cancers. Here, we identified the antibiotic Novobiocin (NVB) as a specific POLθ inhibitor that selectively kills HR-deficient tumor cells *in vitro* and *in vivo*. NVB directly binds to the POLθ ATPase domain, inhibits its ATPase activity, and phenocopies POLθ depletion. BRCA-deficient tumor cells and those with acquired PARPi resistance are sensitive to NVB *in vitro* and *in vivo*. Increased POLθ expression levels predict NVB sensitivity. The mechanism of NVB-mediated cell death in PARPi resistant cells is the accumulation of toxic RAD51 foci, which also provides a pharmacodynamic biomarker for NVB response. Our results demonstrate that NVB may be useful alone or in combination with PARPi in treating HR-deficient tumors, including those with acquired PARPi resistance.

**One Sentence Summary:** We identified Novobiocin as a specific POLθ inhibitor that selectively kills naïve and PARPi resistance HR-deficient tumors *in vitro* and *in vivo.*

---

Since the discovery of the synthetic lethal relationship between PARP inhibition and homologous recombination (HR) repair deficiency (*1*, *2*), PARP inhibitors (PARPi) have become a new line of therapy for HR-deficient cancers (*3–7*). However, resistance to PARPi has emerged as a major obstacle to their clinical effectiveness (*8*). While several mechanisms of PARPi resistance are known (*9*–*14*), a rational and effective method for overcoming such resistance is still lacking. DNA polymerase theta (POLθ or POLQ), a key enzyme in microhomology-mediated end joining (MMEJ), is synthetic lethal with HR and a druggable target in HR-deficient cancers (*15*–*18*). A small-molecule screen identified novobiocin (NVB) as a specific POLQ inhibitor. NVB inhibited POLQ *in vitro* and *in vivo*, Moreover, we demonstrated that NVB can be used alone or in combination with PARPi in treating HR-deficient tumors, even after they have acquired PARPi resistance.

## RESULTS

### A small-molecule screen identified NVB as a specific POLθ inhibitor

Polymerase theta (POLθ, encoded by the *POLQ* gene) is a critical enzyme for repairing DNA double-strand breaks (DSBs) through a microhomology-mediated end joining (MMEJ) mechanism (*19*, *20*). POLθ expression is elevated in several types of cancers where it correlates with poor prognosis (*21*, *22*). HR-deficient tumors upregulate POLθ- mediated MMEJ as a survival strategy, and loss of POLθ in these tumors results in cell death (*15*, *17*). This synthetic lethality with HR defines the rationale for developing POLθ inhibitors for the treatment of HR-deficient cancers (*16*).

The ATPase activity of POLθ is required for the survival of HR-deficient cells (*17*). We took advantage of the strong *in vitro* ATPase activity of purified POLθ (**Fig. S1A**) and designed a large-scale small-molecule screen to identify POLθ inhibitors based on the ADP-Glo luminescent assay (**Fig. 1A, B**). The hits from the primary screen were evaluated in a secondary screen (**Fig. S1B-D**). The four most potent POLθ ATPase inhibitors were Mitoxantrone (MTX), Suramin (SUR), Novobiocin (NVB) and Aurintricarboxylic acid (AUR), which were individually verified in a ^32^P-based radiometric ATPase assay (**Fig. S1E**). To test the specificity of the most potent compounds, we purified two other known DNA repair-related ATPases, namely SMARCAL1 and CHD1 (*23*, *24*), and performed ATPase inhibition assays. Only NVB showed high specific inhibition of POLθ with little effect on SMARCAL1 or CHD1 activity, while MTX, SUR and AUR were nonspecific pan-ATPase inhibitors that inhibited all three ATPases (**Fig. 1C and S1F, G**).

**Figure 1.**
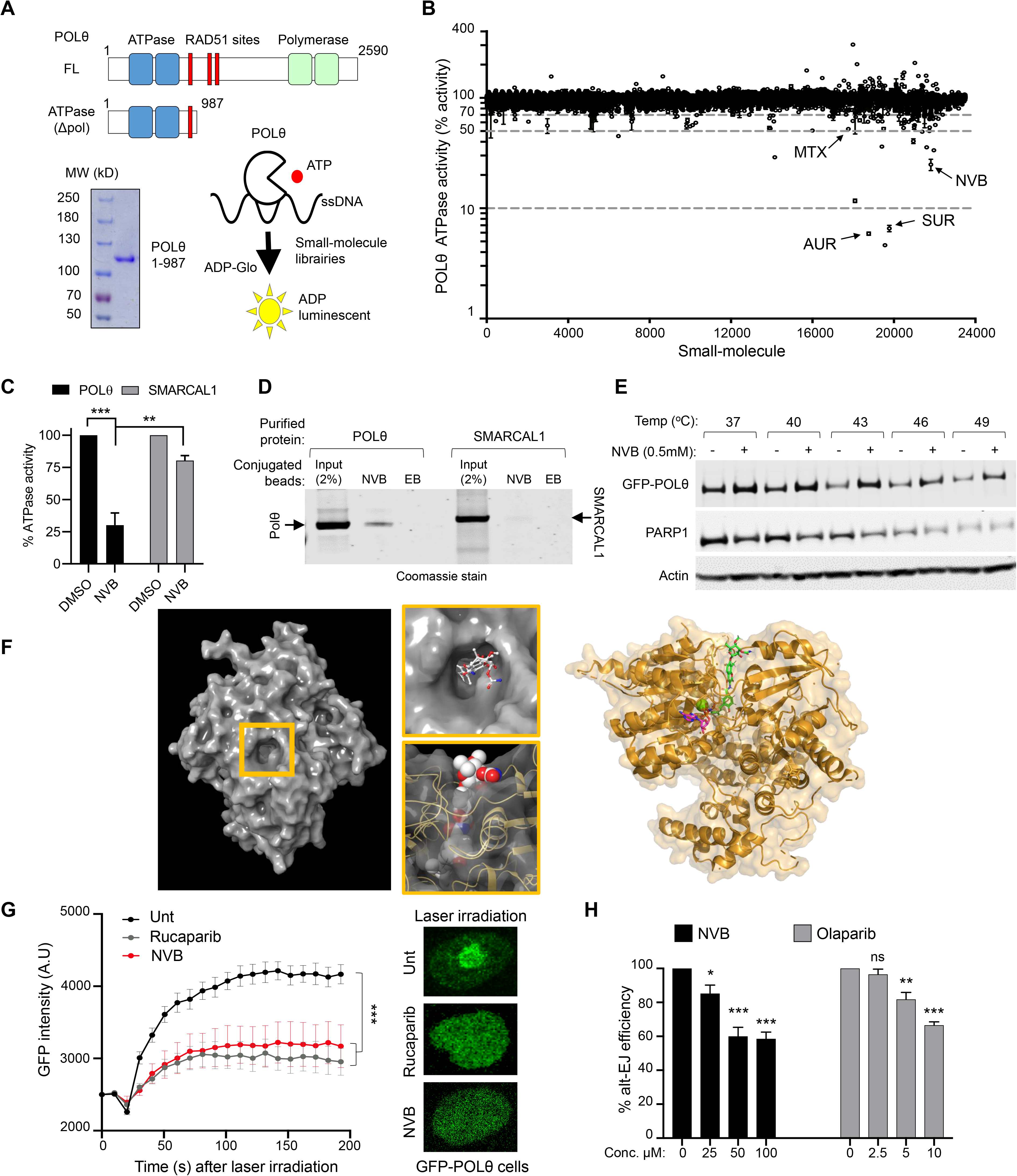
A small-molecule screen identifies Novobiocin (NVB) as a specific POLθ ATPase inhibitor. **A**, The domain structures of full length (FL) and ATPase domain (or ∆Pol, a.a. 1-987) of POLθ, a Coomassie-stained gel of the purified POLθ ATPase domain, and a schematic of the small molecule screen for POLθ inhibitors. **B**, Results of the small-molecule screen. Shown is quantification of POLθ ATPase activity measured by the ADP-Glo assay in the presence of small-molecule libraries. We screened a total of 23,883 small-molecules in duplicate across multiple libraries enriched with known bioactive compounds, and we identified 72 compounds (0.3% of total) that significantly reduced POLθ ATPase activity (*z*-score < −4). Four top hits verified in the secondary screen (Fig. S1B-E) were labeled; Mitoxantrone (MTX), Suramin (SUR), Novobiocin (NVB) and Aurintricarboxylic acid (AUR). Data shown are mean ± s.e.m, n = 2. **C**, Quantification of POLθ and SMARCAL1 ATPase activity in the presence of indicated small-molecules. Mean ± s.d., n = 3. **D**, Pulldown experiments *in vitro* with NVB-conjugated beads and purified POLθ and SMARCAL1 ATPase domains (Flag-tag purification from SF9 cells). Input and pulldown proteins were detected by Coomassie staining on SDS gel. **E**, Thermal shift assay with NVB at indicated temperatures. Full length GFP-POLθ was detected by Western blot using the GFP antibody and PARP1 by a PARP1 antibody. Data in **D** and **E** are representative images of three independent experiments. **F**, The NVB binding tunnel in POLQ ATPase domain (shown in surface - PDB 5AGA) was predicted by extra precision glide docking and lowest binding free energy from prime MM-GBSA calculations to have multiple hydrogen bonds and close hydrophobic packing. Top and side views showing NVB docking into the tunnel. Binding modes of NVB (green sticks) and AMP-PNP (magenta sticks) in POLQ helicase domain (PDB 5AGA) with green sphere showing active site Mg^2+^ ion. **G**, Representative images and quantification of POLθ accumulation at sites of laser micro-irradiated DNA damage in cells overexpressing GFP-tagged, full-length POLθ in the presence of indicated inhibitors. Mean ± s.e.m, n = 3. **H**, MMEJ repair reporter assay in U2OS cells treated with indicated inhibitors. Mean ± s.d., n = 3. Statistics analysis between groups was performed using *t*-test with Welch’s correction. *, *p* < 0.05; **, *p* < 0.01; ***, *p* < 0.001; ns, not significant.

To further evaluate the specificity of Novobiocin (NVB) for POLθ, we analyzed its binding capacity. NVB-conjugated beads pulled down the purified ATPase domain of POLθ but not of SMARCAL1, suggesting that NVB directly binds to the POLθ ATPase domain *in vitro* (**Fig. 1D**, **S1H**). Active NVB-beads also pulled down GFP-tagged full-length POLθ overexpressed in HEK293T cells (**Fig. S1I**). Excess free NVB competed the POLθ binding to the NVB-beads (**Fig. S1J**). Next, we performed protein thermal shift assays, an assay commonly used to demonstrate ligand-protein binding (*25*, *26*). NVB significantly increased the thermal stability of POLθ when compared to DMSO (**Fig. 1E**), while the PARP1 inhibitor olaparib, used as an internal control, effectively stabilized the PARP1 protein (**Fig. S1K**). Finally, we performed molecular docking to find the NVB binding site using the published crystal structure of POLQ ATPase domain (*27*). The computational docking results revealed that a deep tunnel within the POLQ ATPase domain could accommodate NVB, and that one end of NVB could reach the active site to prevent ATP binding to POLQ and inhibit its ATPase activity (**Fig. 1F**). Taken together, these data demonstrate that NVB binds directly and specifically to the POLθ ATPase domain *in vitro*.

We next assessed whether cellular exposure to NVB has the same consequences as POLθ depletion. Both the ATPase and the polymerase domains contribute to efficient MMEJ repair activity (*28*), and the PARP-dependent recruitment of POLθ to DSB sites is a crucial step in this function (*15*, *17*). The inhibition of POLθ by NVB prevented the recruitment of POLθ to laser micro-irradiated DNA damage sites in human cells (**Fig. 1G**). Consequently, NVB inhibited the MMEJ activity in U2OS cells, as demonstrated by the GFP-based EJ2 repair assay (**Fig. 1H**). In contrast, NVB had little effect on HR activity, as measured by the DR-GFP reporter assay (**Fig. S2A**). POLθ also functions as an anti-recombinase that antagonizes HR, and depletion of POLθ increases RAD51 (*17*) and γH2AX (*29*, *30*) foci assembly. Again, cellular exposure to NVB recapitulated the RAD51 and γH2AX phenotypes of a POLθ knockdown. NVB activated more RAD51 foci (**Fig. S2B**) and γH2AX foci (**Fig. S2C**) in RPE1 cells, regardless of IR exposure. Taken together, these results establish that NVB-mediated POLθ inhibition impairs POLθ DNA repair function and phenocopies POLθ depletion in human cells.

### NVB specifically Kills HR-deficient cells *in vivo* and *in vitro*

Depletion of POLθ is synthetically lethal in HR-deficient tumor cells (*15*, *17*). Since NVB has been previously approved for human use, we decided to directly test whether NVB reduces the survival of HR-deficient tumor cells in animal models. First, we evaluated the efficacy of NVB in tumors derived from the K14-Cre-*Brca1*^*f/f*^*;Trp53*^*f/f*^ genetically-engineered mouse model (GEMM) of triple-negative breast cancer (TNBC), in which spontaneous mammary carcinomas develop after approximately 7 months (*31*, *32*). Tumors from this model were transplanted into immunocompetent FVB/129P2 syngeneic mice that were subsequently treated with NVB. NVB strongly suppressed the growth of these *Brca1*- deficient GEMM-derived tumors (**Fig. 2A** and individual mouse data in **Fig. S3A**). The NVB-treated tumors were significantly smaller compared to the vehicle-treated after 7 days of treatment and beyond. NVB prolonged the overall survival of the tumor-bearing mice by 3-fold compared to vehicle, with median survival times of 29 and 10 days for NVB- and vehicle-treated mice, respectively (**Fig. 2B**).

**Figure 2.**
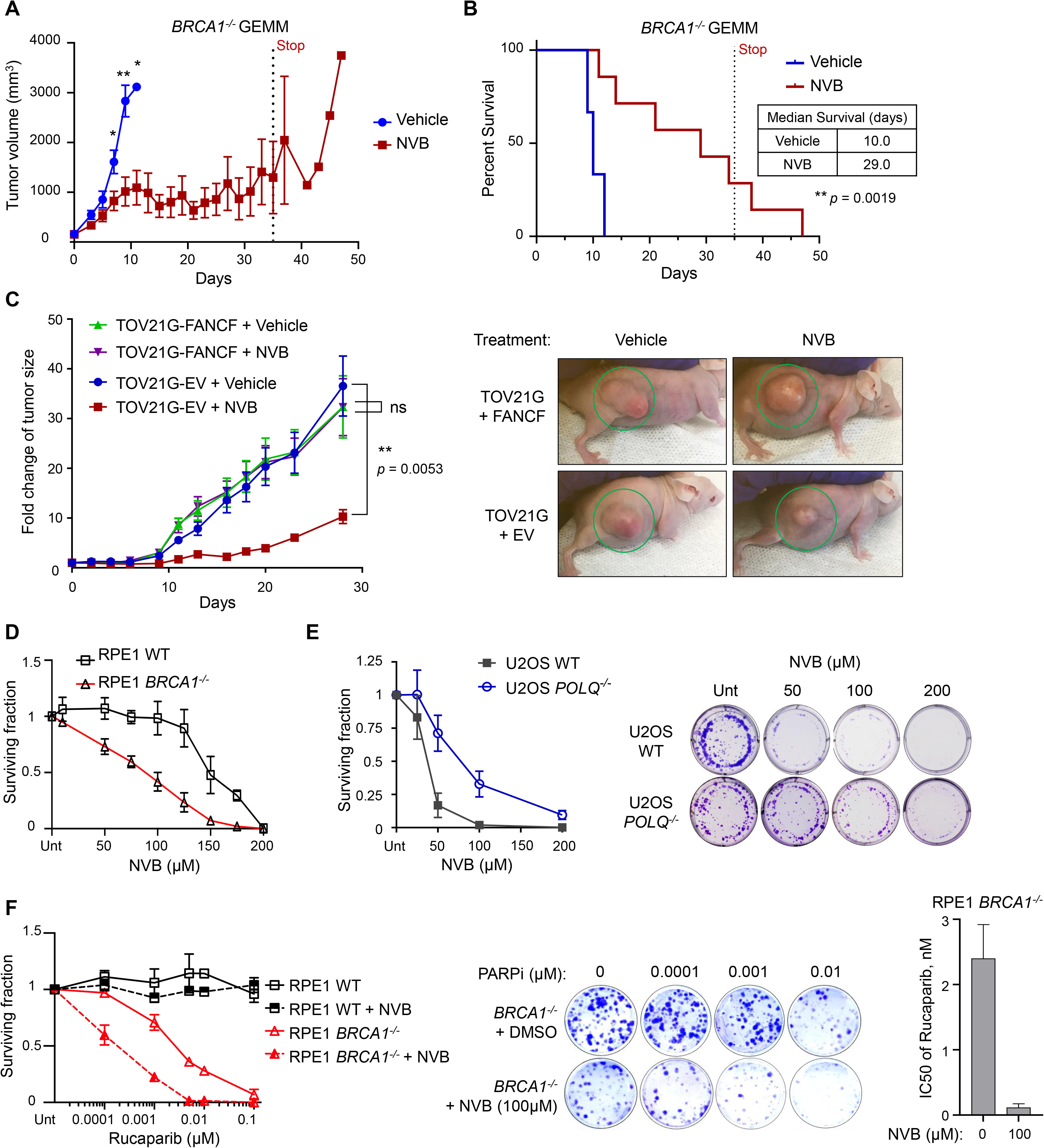
NVB kills HR-deficient tumors *in vivo* and *in vitro* and synergizes with PARPi. **A**, Tumor growth of the GEMM model (*Brca1*^*−/−*^ TNBC) after treatment with vehicle (PBS) or NVB. Tumor chunks from GEMM mice were implanted in syngeneic FVB/129P2 mice, which were treated with PBS or 100 mg/kg NVB via IP injection twice a day for 5 weeks. n = 6 for PBS group and n = 7 for NVB group. Mean ± s.e.m. is shown. Statistical analyses were *t*-test. **B**, Survival plot of the experiment shown in (**A**). Median survival and *p* value are shown. Statistics analyses were *t*-test. **C**, Tumor growth of the xenograft model with TOV21G (FANCF deficient ovarian cancer) or FANCF-complemented TOV21G cells. The tumor bearing mice were treated with vehicle or 100 mg/kg NVB via IP injection twice a day for 4 weeks. Fold change of tumor was calculated using the formula Fold = T/T0, where T is the tumor size at given time and T0 is the initial tumor size. Data are Mean ± s.e.m., n = 10 in each group. Dunn’s multiple comparisons test. Representative images are TOV21G xenograft tumors at the end of NVB treatment. **D,** Clonogenic survival of *Brca1*^*−/−*^ and WT RPE1 cells under increasing concentrations of the POLθ inhibitor NVB. Survival is normalized to the untreated (Unt) sample. Data shown are mean ± s.e.m., n = 4. **E**, Representative images and quantification of clonogenic survival of *POLQ*^*−/−*^ and WT U2OS cells under increasing concentrations of NVB. Survival is shown as relative to the untreated (Unt) sample. Data shown are mean ± SD, n = 5. **F,** Clonogenic survival of *Brca1*^*−/−*^ and WT RPE1 cells under increasing concentrations of the PARPi Rucaparib alone or in combination with NVB. Quantifications, representative images and calculated IC50 values are shown. Data shown are mean ± s.e.m., n = 3.

Next we tested whether the efficacy of NVB *in vivo* was specific to HR-deficient tumors. We performed a mouse xenograft study with a FANCF-deficient, HR-deficient ovarian cancer cell line [TOV21G + empty vector (EV)] or with the FANCF-complemented, HR-proficient cell line (TOV21G + FANCF cDNA) (**Fig. 2C**). TOV21G + EV cells but not TOV21G + FANCF cDNA cells were sensitive to olaparib and NVB (**Fig. S3B, C**) (*33*). While vehicle treatment had no impact on the growth of FANCF-deficient or FANCF-complemented tumors, NVB specifically impaired the growth of the FANCF-deficient tumors, with no effect on FANCF-complemented tumors (**Fig. 2C**). We also measured RAD51 foci as a pharmacodynamic biomarker and showed that RAD51 foci were strongly induced in the treated tumors (**Fig. S3D-F**). These results demonstrate that NVB strongly and specifically suppresses the growth of HR-deficient tumors *in vivo*.

To confirm and strengthen the *in vivo* results, we studied isogenic pairs of HR-deficient and HR-proficient cells *in vitro*. We generated *BRCA1* (*34*) and *BRCA2* knockout RPE1 cells (referred as *BRCA1*^−/−^ and *BRCA2*^*−/−*^) in a *TP53*^−/−^ background, which were sensitive to PARPi (**Fig. S4A**). Consistent with our *in vivo* data, clonogenic survival assays showed that NVB significantly reduced the survival of *Brca1*^*−/−*^ and *BRCA2*^*−/−*^ cells, compared to the isogenic WT cells (**Fig. 2D**, **S4B and Fig. S4C**). We also tested the top 15 initial hits from our screen for their ability to differentially affect the survival of *Brca1*^*−/−*^ and WT cells. Among those, NVB had the highest efficacy in selectively killing the *Brca1*^*−/−*^ cells (**Fig. S4D**).

To demonstrate the cellular outcome of NVB exposure, we next showed that NVB exposure induces apoptosis in *Brca1*^*−/−*^ but not WT cells in a dose dependent manner (**Fig. S5A**). NVB induced chromosomal aberrations and radial chromosomes (a marker of genomic instability) in *Brca1*^*−/−*^ cells, even in the absence of the crosslinking agent mitomycin C (MMC) (**Fig. S5B-D**), confirming that NVB increases DNA damage. These results further demonstrate that NVB phenocopies POLθ depletion, and indicate that NVB-mediated POLθ inhibition is synthetically lethal with HR deficiency and specifically induces cell death in HR-deficient tumor cells.

### POLθ is the major target of NVB in human cells

Off target effects are major concerns of biological small inhibitors. To further evaluate whether POLθ is the specific target of NVB and to demonstrate the specificity of NVB in cells, we knocked out the *POLQ* gene in U2OS and RPE1 cells using CRISPR-Cas9 and asked whether loss of POLθ alleviates the cytotoxicity of NVB (**Fig. 2E and S6**). POLQ knockout cells were confirmed by genomic sequencing, immunoblotting, increased RAD51 foci, increased sensitivity to irradiation, and increased micronuclei formation (**Fig. S6A-F**). Importantly, *POLQ*^*−/−*^ cells showed increased tolerance to NVB, both in U2OS and RPE1 cells, as compared to the WT cells (**Fig. 2E and S6G**), indicating that POLθ is the major target of NVB in human cells.

NVB is a known coumarin antibiotic that inhibits DNA gyrase in bacteria (*35*). NVB has been reported to inhibit the ATPase of HSP90 and of the DNA gyrase homolog topoisomerase II (TOP2) in eukaryotes (*36*) (*37*) (*38*), suggesting that these enzymes may be off-target enzymes of NVB. However, other reports suggest that NVB does not inhibit TOP2 in eukaryotic cells (*39*). Additionally, the reported IC_50_ of these targets are very high (approximately 700 μM for HSP90 and 300 μM for TOP2), compared with the IC_50_ of POLθ. Nevertheless, we next examined whether the cytotoxic effect of NVB in HR-deficient cells resulted from off-target inhibition of HSP90 or TOP2. Unlike the well-characterized HSP90 inhibitor PU-H71, NVB treatment at a concentration that killed HR-deficient cells (i.e. 100 μM) did not induce the degradation of HSP90 clients such as AKT1, BRCA1, or CDK6, or increase the expression of HSP70 (**Fig. S7A-C**). These results suggest that the cytotoxic effect of NVB is not related to HSP90 inhibition. Furthermore, the combination of NVB and the TOP2 inhibitor etoposide exhibited additive cytotoxicity in TOV21G and CAPAN1 cells (**Fig. S7D, E**), suggesting that NVB and etoposide do not share TOP2 as a target and that NVB cytotoxicity is unrelated to TOP2 inhibition.

### NVB potentiates the effect of PARPi in HR-deficient tumors

PARPi are currently being evaluated in different combination therapy settings with cytotoxic chemotherapy, radiation, targeted therapies, and immunotherapies (*40*). We explored whether the combination of NVB and a PARPi may be more effective than PARP inhibition alone in killing HR-deficient tumors. Indeed, NVB further sensitized *BRCA1*^−/−^ RPE1 cells to the PARPi rucaparib, but not BRCA-proficient WT cells in clonogenic assays (**Fig. 2F**). We studied the combination with a second PARPi, olaparib, where synergism of NVB and olaparib was also observed in HR-deficient TOV21G+EV cells, but not in HR-proficient TOV21G+FANCF cells (**Fig. S7F**). Importantly, NVB reduced the IC_50_ of rucaparib by more than 20-fold in *BRCA1*^−/−^ RPE1 cells and the IC_50_ of olaparib more than 40-fold in TOV21G cells compared to their respective HR-proficient counterparts (**Fig. 2F and S7F**). One strategy to prevent acquired resistance during treatment is combination therapy. The observed synergy between NVB and PARPi prompted us to evaluate whether NVB could overcome PARPi resistance in HR-deficient tumors.

### POLθ inhibition by NVB overcomes acquired PARPi resistance *in vivo* and *in vitro*

Multiple PARP inhibitors (PARPi) including olaparib, niraparib, rucaparib, and talazoparib have received approval for the treatment of ovarian and breast tumors with HR deficiency (*41*–*43*). Despite the remarkable initial tumor response and improved progression-free survival (PFS) in HR-deficient patients (*4*–*6*, *44*), acquired PARPi resistance is emerging as an unmet medical need (*8*).

To evaluate whether POLθ inhibition could represent a novel therapeutic option for PARPi-resistant tumors, we first tested the efficacy *in vivo* of NVB in the patient-derived xenograft (PDX) model DF83, generated from an HR-deficient (loss of RAD51C expression), human high-grade serous ovarian carcinoma (*45*). NSG mice bearing the HR-deficient DF83 ovarian cancer PDX model were treated with olaparib, NVB, or a combination of the two, and tumor growth was monitored by bioluminescence imaging (BLI) (**Fig. 3A**). In order to determine the combined effect with NVB, olaparib was delivered at a submaximal dose (50 mg/kg daily), and we observed only a small degree of tumor growth inhibition. Treatment of DF83 with NVB alone (75 mg/kg, twice a day) led to tumor regression, while tumors in vehicle-treated mice showed exponential growth (**Fig. 3B**). Strikingly, when the DF83 tumor-bearing mice were given NVB and olaparib in combination, we observed complete tumor regression, and few tumor cells were detectable via BLI on day 28 in this group (**Fig. 3A-B**). These *in vivo* results suggest NVB may be useful in combination with PARPi to achieve better efficacy and prevent drug resistance in HR-deficient tumors. No significant toxicity to normal mouse tissue was observed from the drug combination.

**Figure 3.**
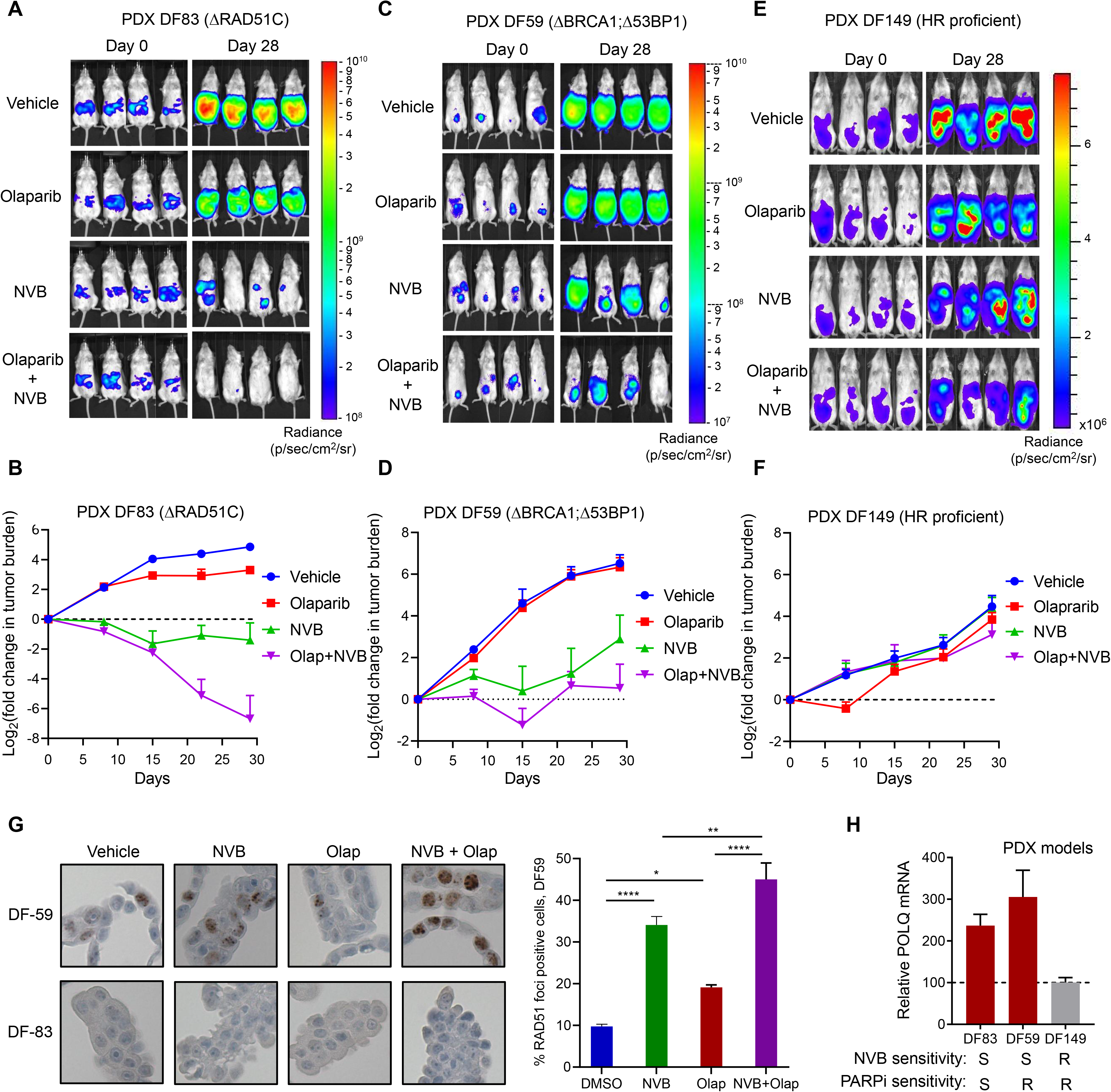
NVB overrides PARPi resistance *in vivo* in PDX models and RAD51 is a dynamic pharmacodynamic biomarker. **A-B**, Efficacy of NVB in PDX model DF83 (RAD51C deficient). NSG mice (n = 7 mice/group) bearing luciferized DF83 cells derived from an ovarian cancer patient were treated with 50 mg/kg of olaparib (daily, orally), 75 mg/kg NVB (via IP, b.i.d.), or both for 4 weeks. Tumor growth was monitored weekly by bioluminescence imaging. Representative images of tumor burden on day 0 and day 28 are shown, and quantifications shown are mean ± s.e.m. Log_2_(fold change in tumor size) was calculated using the formula log_2_(T/T0), where T is the tumor volume at given time and T0 is the initial tumor volume. **C-D**, Efficacy of NVB in PARPi resistant PDX model DF59 (BRCA1-deficient, 53BP1 loss), n = 6 mice/group. Same conditions and analysis were applied as in **A-B**. **E-F**, Efficacy of NVB in PARPi resistant PDX model DF149 (BRCA1-WT, HR proficient), n = 7 mice/group. Same conditions and analysis were applied as in **A-B**. **G**, Representative IHC images and quantifications of RAD51 foci in DF-59 and DF-83 PDX models. Tumor bearing mice were dosed with indicated drugs and as in efficacy study in Fig 3, and tumor cells were collected from ascites 52 hours after first dose. FFPE sections of the isolated tumor cells were stained with a RAD51 antibody. The percentage of RAD51-foci positive cells (> 4 foci/cell) in 3 random 40X fields was estimated in DF-59. No RAD51 focus was observed in the DF-83 model. Statistical analysis method was one-way ANOVA Bonferroni’s multiple comparisons test. *, *p* < 0.05; **, *p* < 0.01; ****, *p* < 0.0001. **H**. POLQ expression at mRNA level in PDX models DF53, DF83 and DF149.

Next, we evaluated the efficacy of NVB alone and in combination with olaparib in the PARPi-resistant PDX model DF59, which was isolated from a patient with a germline *BRCA1* mutation and acquired PARPi resistance. This model harbors biallelic mutations in the *TP53BP1* locus, providing a mechanism of the acquired PARPi resistance (*45*). In the DF59 PDX model, there was no response to olaparib monotherapy, but NVB monotherapy substantially reduced tumor growth, indicating a lack of cross-resistance between PARP inhibition and NVB. Importantly, the combination of NVB and olaparib further inhibited DF59 tumor growth, with tumor regression in the first two weeks (**Fig. 3C-D**). In contrast to DF83 and DF59 models, the BRCA1 WT, HR proficient PDX model DF149 was resistant to both drugs and to the combination (**Fig. 3E-F**). These data demonstrate that NVB can inhibit the growth of a *BRCA1*-mutated, PARPi-resistant tumors *in vivo* and that the combination of NVB and a PARPi is particularly effective in treating at least some PARPi-resistant tumors.

### RAD51 focus is a pharmacodynamic biomarker for NVB response *in vivo*

We previously showed that depletion of POLθ in HR-deficient tumors resulted in RAD51 accumulation (*17*). Similarly, 53BP1 and POLθ are synthetic lethal, and the double knockout cells accumulate large, toxic RAD51 foci (*46*). We thus reasoned that NVB may induce RAD51 foci in HR-deficient tumors with PARPi resistance *in vivo*, and the RAD51 foci may serve as a pharmacodynamic biomarker for NVB response. To test this hypothesis, we collected tumor cells from the PARPi and/or NVB treated PDX models and evaluated RAD51 foci by immunohistochemistry (**Fig. 3G)**. Since DF83 is an HR-deficient and PARPi sensitive model, RAD51 foci were not observed. The HR-restored and PARPi-resistant DF59 model, with biallelic 53BP1 mutations, exhibited a strong increase in RAD51 foci when treated with NVB alone or in combination with olaparib (**Fig. 3G)**. Similarly, TOV21G xenograft tumors also exhibited increased RAD51 foci by immunohistochemistry *in vivo* after NVB treatment, which was further enhanced by olaparib (**Fig. S3D-F**). Results from these two *in vivo* studies demonstrate that RAD51 focus can serve as a pharmacodynamic biomarker for the response of tumors to NVB or other POLθ inhibitors. Interestingly, we also observed that two NVB-responsive tumor models had elevated levels of POLθ mRNA expression, compared to the NVB-resistant DF149 model, suggesting that elevated POLθ may be a useful predictive biomarker for POLθ inhibitor response (**Fig. 3H**).

### NVB overcomes multiple PARPi resistance mechanisms but not perfect BRCA reversions

To better understand the mechanism by which NVB overcomes PARPi resistance and, we generated several PARPi-resistant clones from *BRCA1*^*−/−*^ *RPE1* cells by gradually exposing them to increasing concentrations of PARPi (**Fig. 4A**). We selected four clones (named R1 to R4), all of which have acquired resistance to olaparib (**Fig. S8A-C**). Multiple PARPi resistance mechanisms were identified in these clones, corresponding to published mechanisms (*8*). Replication fork stabilization was evident in the R1 and R2 clones (**Fig. S8D**). The R1, R3, and R4 clones have restored RAD51 foci, a marker for restored HR activity (**Fig. S8E**). The restoration of HR repair in these clones may have resulted in part from their downregulation of NHEJ repair. Indeed, the R3 clone had decreased REV7 expression, and the R4 clone had decreased 53BP1 expression (**Fig.S8F-G**). None of the clones had re-expressed BRCA1, thereby excluding somatic reversion of BRCA1 as an underlying PARPi resistance mechanism (**Fig. 4B and S8H**). Strikingly, all four PARPi-resistant clones remained sensitive to NVB to a similar extent as the parental RPE-*BRCA1*^−/−^ cells (**Fig. 4C-D**), suggesting that NVB may overcome multiple mechanisms of acquired resistance to PARPi. To confirm that POLθ inhibition was responsible for cell death in the PARPi-resistant *BRCA1*^−/−^ clones treated with NVB, we knocked out *POLQ* in R1, R2, and the parental cells and measured their cell survival. Upon POLQ knockout, the PARPi-resistant clones showed reduced survival similar to the parental *BRCA1*^−/−^ cells, demonstrating that POLQ inhibition was the mechanism for NVB cytotoxicity in these cells (**Fig. S8I**).

**Figure 4.**
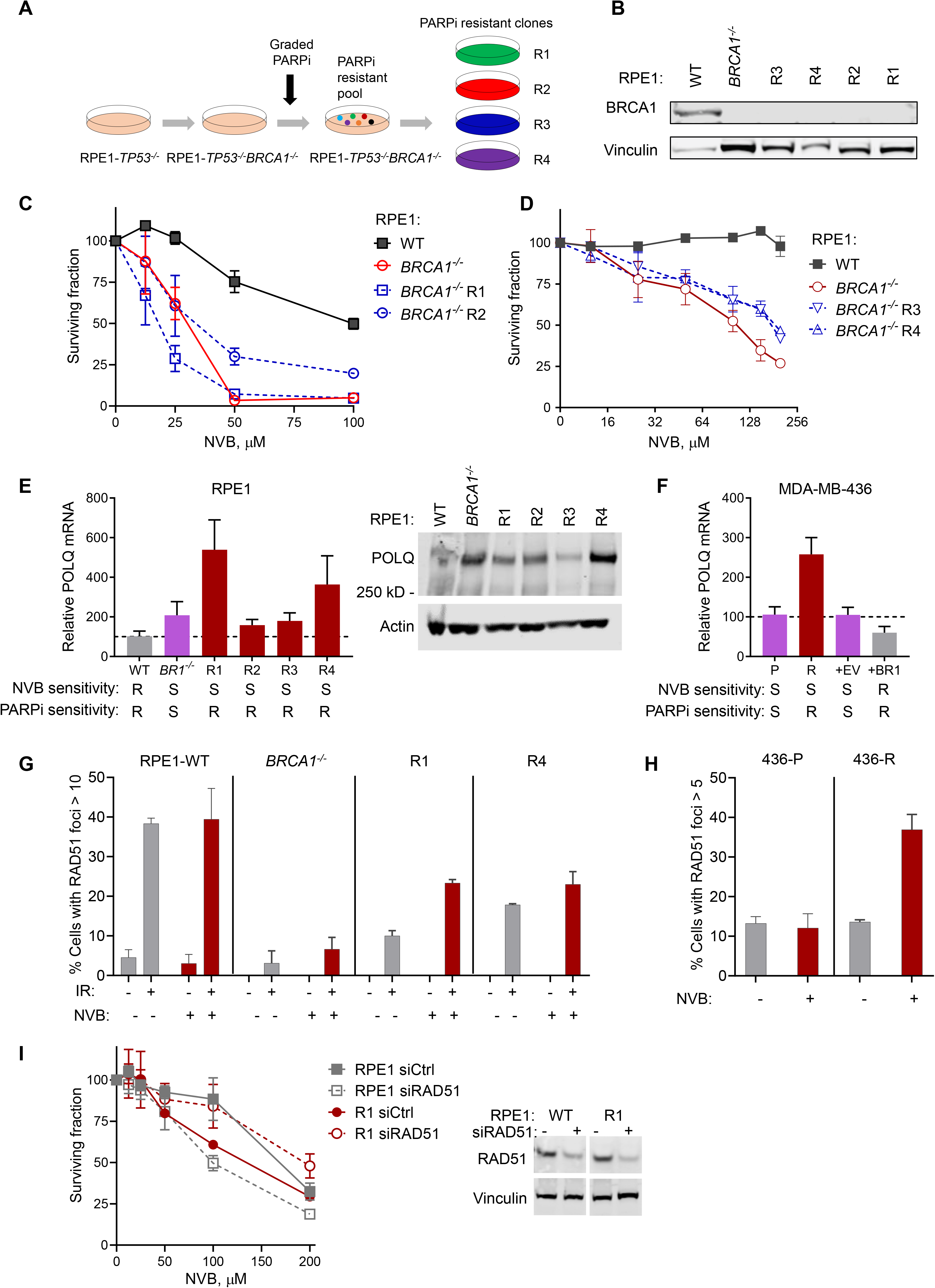
POLQ expression level is a predictive biomarker for NVB sensitivity. **A**. A schematic shows how the PARPi resistant clones R1-R4 were generated. TP53 was knocked out in order to generate *BRCA1* knockout (*Brca1*^*−/−*^) RPE1 cells, which were then used for selection of PARPi resistant clones. **B**. A Western blot of WT and *Brca1*^*−/−*^ RPE1 cells and the PARPi resistant clones, using an anti-BRCA1 antibody (Millipore #OP92). No BRCA1 reversion was observed in the clones. **C-D.** NVB sensitivity of WT RPE1, *Brca1*^*−/−*^ RPE1, and PARPi-resistant *Brca1*^*−/−*^ clones (R1-R4). R1 was selected by olaparib and R2-R4 were selected by Niraparib. R1 and R2 were tested in clonogenic survival assay **(C)**. R3 and R4 were tested in CellTiter-Glo cell viability assay (**D**). Data are mean ± s.e.m., n = 3. **E**. POLQ expression at mRNA level and protein level in WT RPE1, *Brca1*^*−/−*^ RPE1, and PARPi-resistant *Brca1*^*−/−*^ clones (R1-R4). POLQ mRNA was measured by qRT-PCR and normalized to beta-Actin (ACTB). **F**. POLQ expression at mRNA level in parental (P), PARPi resistant (R), and empty vector (+EV) or BRCA1-cDNA (+BR1) complemented MDA-MB-436 cells. **G**, RAD51 foci assay with DMSO or NVB in RPE1-*BRCA1* WT RPE1, *Brca1*^*−/−*^ RPE1, and PARPi-resistant *Brca1*^*−/−*^ clones (R1-R4) 4 hours after 5 Gy of IR. Cells with more than 10 foci were counted as positive. Data were Mean ± SEM from two independent experiments. **H**, RAD51 foci assay with DMSO or NVB in MDA-MB-436 or the PARPi-resistant MDA-MB-436-R. Cells with more than 5 foci were counted as positive. Data were Mean ± SEM from two independent experiments. **I.** Novobiocin sensitivity of RPE1-WT and the R1 PARPi-resistant clone after being treated with Control or RAD51 siRNA. A Western blot shows the efficiency of RAD51 knockdown in RPE1-WT and the R1 PARPi-resistant clone.

Next, we examined more clinically relevant PARPi-resistant, HR-deficient models, using tumor cells derived from patients. First, we evaluated the effect of NVB on a PARPi resistant clone (MDA-MB-436-R), previously derived from the BRCA1-mutated breast cancer cell line MDA-MB-436 (*47*). These resistant cells restored HR by two mechanism-namely, stabilization and increased expression of a mutant BRCA1 protein and loss of 53BP1-resulting in increased RAD51 loading. While highly resistant to olaparib, the MDA-MB-436-R cells were NVB sensitive, like the parental cells (**Fig. S9A-B**). In addition, we studied the PARPi-resistant clone UWB1.289-YSR12 (derived from the *BRCA1*-null human ovarian cancer cell line UWB1.289), which acquired resistance to PARPi via stabilization of replication forks (*48*). Although highly resistant to olaparib, these cells also remained sensitive to NVB, like the parental cells (**Fig. S9C-D**), once again suggesting that acquired resistance to PARPi does not determine cross-resistance to NVB.

Another known mechanism of PARPi resistance in *BRCA1/2*^*−/−*^ tumors in the clinic is somatic reversion of the BRCA1/2 genes (*18*). We next determined whether NVB could overcome PARPi resistance caused by BRCA2 reversion. Unlike other PARPi resistant cells we analyzed, BRCA2-deficient tumor cells with near-perfect somatic reversion or CRISPR-edited reversion of BRCA2 (PEO4 and CAPAN1-CR) (*49*, *50*) were not only resistant to PARPi but also to NVB, when compared to their HR-deficient parental cell lines (PEO1 and CAPAN1) **(Fig. S10A-D)**. These data suggest that NVB can overcome some but not all mechanisms of PARPi resistance.

### POLθ expression is a predictive biomarker for NVB sensitivity

POLθ mRNA expression is increased in HR-deficient tumors and in several other cancers associated with poor prognosis (*17*, *21*). We asked whether increased POLθ mRNA and protein expression may serve as a predictive biomarker for NVB sensitivity. As expected, CRISPR-mediated knockout of *BRCA1* in RPE-1 cells resulted in increased POLθ expression as compare to the parental cells (**Fig. 4E**). Interestingly, all PARPi-resistant clones (R1-R4) retain high POLθ expression, revealing a direct correlation between POLθ expression and NVB sensitivity (**Fig. 4E**). This correlation was also present in the BRCA1-deficient MDA-MB-436 cells. MDA-MB-436-R cells with acquired PARPi-resistance exhibited elevated POLθ mRNA expression and retained NVB sensitivity, while complementation of the parental cells with WT BRCA1 cDNA decreased POLθ expression and NVB sensitivity (**Fig. 4F and S9A-B**). More importantly, the direct correlation of POLθ expression and NVB sensitivity was observed in HR-deficient PDX models, where the elevated POLθ expression in DF83 and DF59 correlated with their high sensitivity to NVB *in vivo* (**Fig. 3H and 3A-F**).

Conversely, low POLθ expression was a predictive biomarker for NVB resistance. In the HR-proficient PDX cells (DF149) with wildtype BRCA1 and in the PARPi-resistant cell lines with BRCA2 reversion mutations (PEO4 and CAPAN1-CR), the cellular resistance to NVB was associated with low POLθ expression levels (**Fig. 3G and S10E**). Also, the loss of POLθ in *POLQ*^*−/−*^ cells correlated with resistance to NVB (**Fig. 2E**). Taken together, in all cases examined, POLθ expression level correlated with cellular sensitivity to NVB, suggesting that POLθ expression in a tumor biopsy could provide a convenient predictive biomarker for patient enrollment in future clinical trials with NVB or other POLθ inhibitors.

### Toxic RAD51 foci is a mechanism of NVB cytotoxicity in PARPi resistant cells

Depletion of POLθ in HR-deficient tumors resulted in RAD51 accumulation (*17*), which can be toxic if not regulated. Similarly, 53BP1 and POLθ are synthetic lethal, and the double knockout cells accumulate toxic RAD51 foci (*46*). We reasoned that accumulation of toxic RAD51 foci induced by NVB may be responsible for killing the PARPi resistant tumor cells. This is supported by the RAD51 pharmacodynamic results, where RAD51 foci were greatly increased in tumors after NVB treatment *in vivo* (**Fig. 3G and S3D-F**). We next determined RAD51 foci dynamics in our PARPi resistant cell line models. NVB induced accumulation RAD51 foci in PARPi-resistant RPE1-BRCA1^−/−^ clones (R1 and R4) more readily than in WT RPE1 cells after irradiation (**Fig. 4G**). Similarly, NVB treatment alone strongly induced RAD51 foci in the PARPi resistant MDA-MB-436-R cells (**Fig. 4H**). In addition, knockdown of RAD51 sensitized RPE1 WT cells to NVB but rescued *Brca1*^*−/−*^ R1 clone from NVB cytotoxicity (**Fig. 4I**). Taken together, these results suggest toxic RAD51 foci, induced by NVB, is a mechanism for NVB cytotoxicity in HR-deficient cells with acquired PARPi resistance.

## DISCUSSION

In summary, NVB is a specific and potent POLθ inhibitor which phenocopies POLθ depletion and is synthetic lethal in HR-deficient tumors. Many tumors types with PARPi resistance remain sensitive to NVB, depending on their mechanism of resistance. Tumors which acquire PARPi resistance by downregulating NHEJ and partially restoring HR repair remain sensitive to NVB. For these tumors, the persistent high level of POLθ expression is a predictive biomarker for their NVB responsiveness, and the NVB-induced RAD51 accumulation is both a pharmacodynamic biomarker and a mechanism of NVB cytotoxic activity.

Finally, our findings may translate to a potential new therapy for treating HR-deficient tumors in addition to PARP inhibition. Importantly, NVB, which is an oral and well-tolerated drug, has previously been investigated in a Phase 1 trial in which partial responses were documented, even though the enrolled patients were not preselected to have HR-deficient cancers (*51*). Results in this study provide a strong rationale for future clinical studies of NVB alone or in combination with PARPi to prevent or overcome PARPi resistance.

## ACKNOWLEDGEMENTS

We thank the ICCB-Longwood Screening Facility at Harvard Medical School, for their help in small-molecule screening. We thank Prafulla Gokhale and Qing Zeng at Belfer Center for Applied Cancer Science for their help in PDX studies. We thank Connor Clairmont for providing RPE *Brca1*^*−/−*^ cells and Lisa Moreau for her help in chromosome aberration assays. We thank Drs. Huy Nguyen and Sandor Spisak for genomic analysis of the *POLθ* Knockout clones. We thank Dr. Jeremy Stark at Beckman Research Institute of the City of Hope for providing DR-GFP and EJ-2 repair substrates.

## DATA AND MATERIALS AVAILABILITY

All related data have been presented in the main manuscript or in supplementary data.

## SUPPLEMENTARY MATERIALS

Figs. S1 to S10

Legends for Figs. S1 to S10

Materials and Methods

Supplementary References

