## SUPPLEMENTARY MATERIALS for "Polymerase Theta Inhibition Kills Homologous Recombination Deficient Tumors"

**This PDF file includes:**

Figs. S1 to S10
Legends for Figs. S1 to S10
Materials and Methods
Supplementary References

#### Fig S1

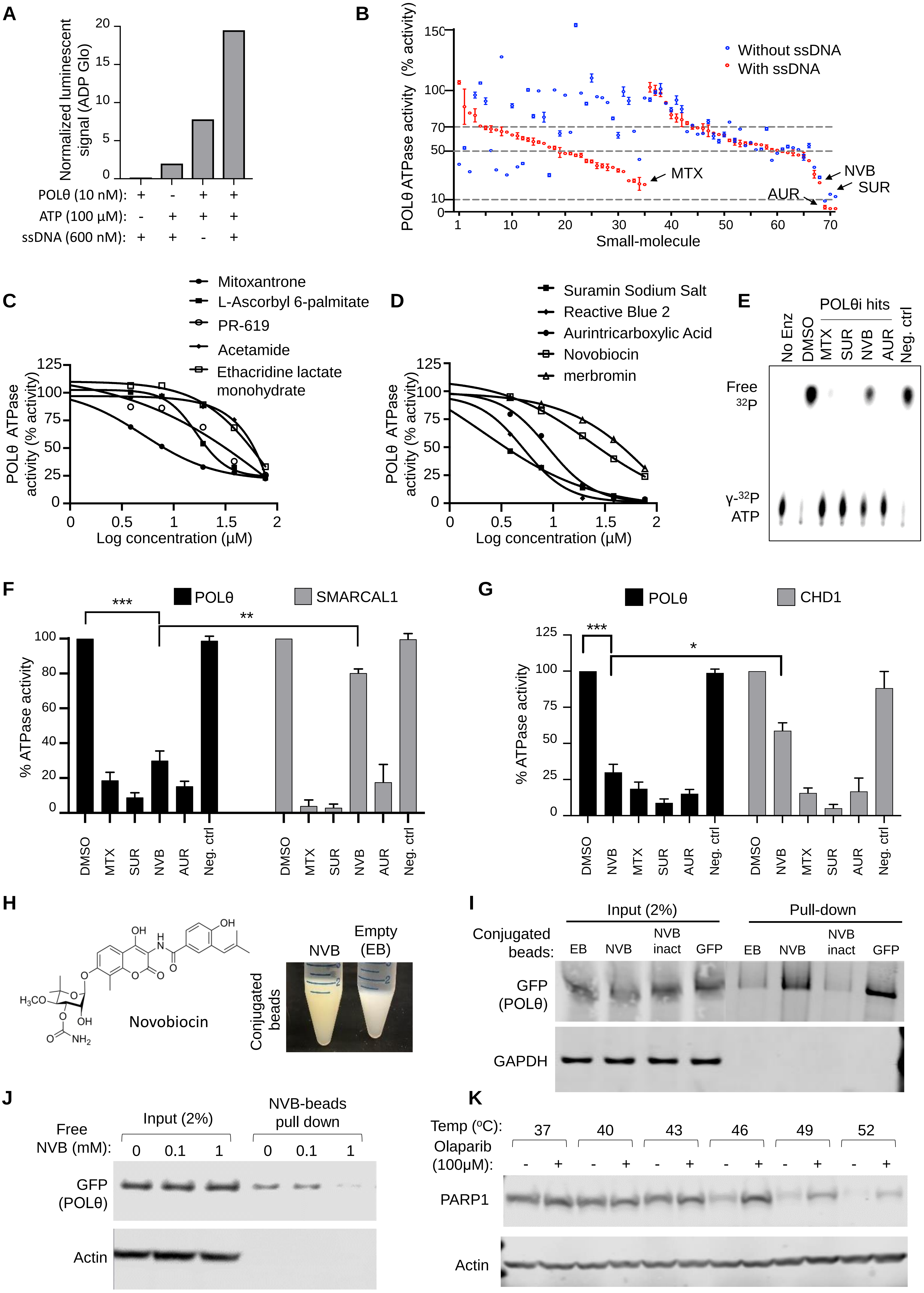

A
Normalized luminescent signal (ADP Glo)
POLθ (10 nM):
ATP (100 µM):
ssDNA (600 nM):
B
POLθ ATPase activity (% activity)
Without ssDNA
With ssDNA
MTX
NVB
SUR
AUR
Small-molecule
C
Mitoxantrone
L-Ascorbyl 6-palmitate
PR-619
Acetamide
Ethacridine lactate monohydrate
POLθ ATPase activity (% activity)
Log concentration (µM)
D
Suramin Sodium Salt
Reactive Blue 2
Aurintricarboxylic Acid
Novobiocin
merbromin
E
POLθi hits
No Enz
DMSO
MTX
SUR
NVB
AUR
Neg. ctrl
Free ³²P
γ-³²P ATP
F
POLθ
SMARCAL1
% ATPase activity
G
POLθ
CHD1
H
Novobiocin
NVB
Empty (EB)
Conjugated beads
I
Input (2%)
Pull-down
Conjugated beads:
EB
NVB
NVB inact
GFP
GFP (POLθ)
GAPDH
J
Input (2%)
NVB-beads pull down
Free NVB (mM):
Actin
K
Temp (°C):
Olaparib (100µM):
PARP1
Actin

#### Fig S2

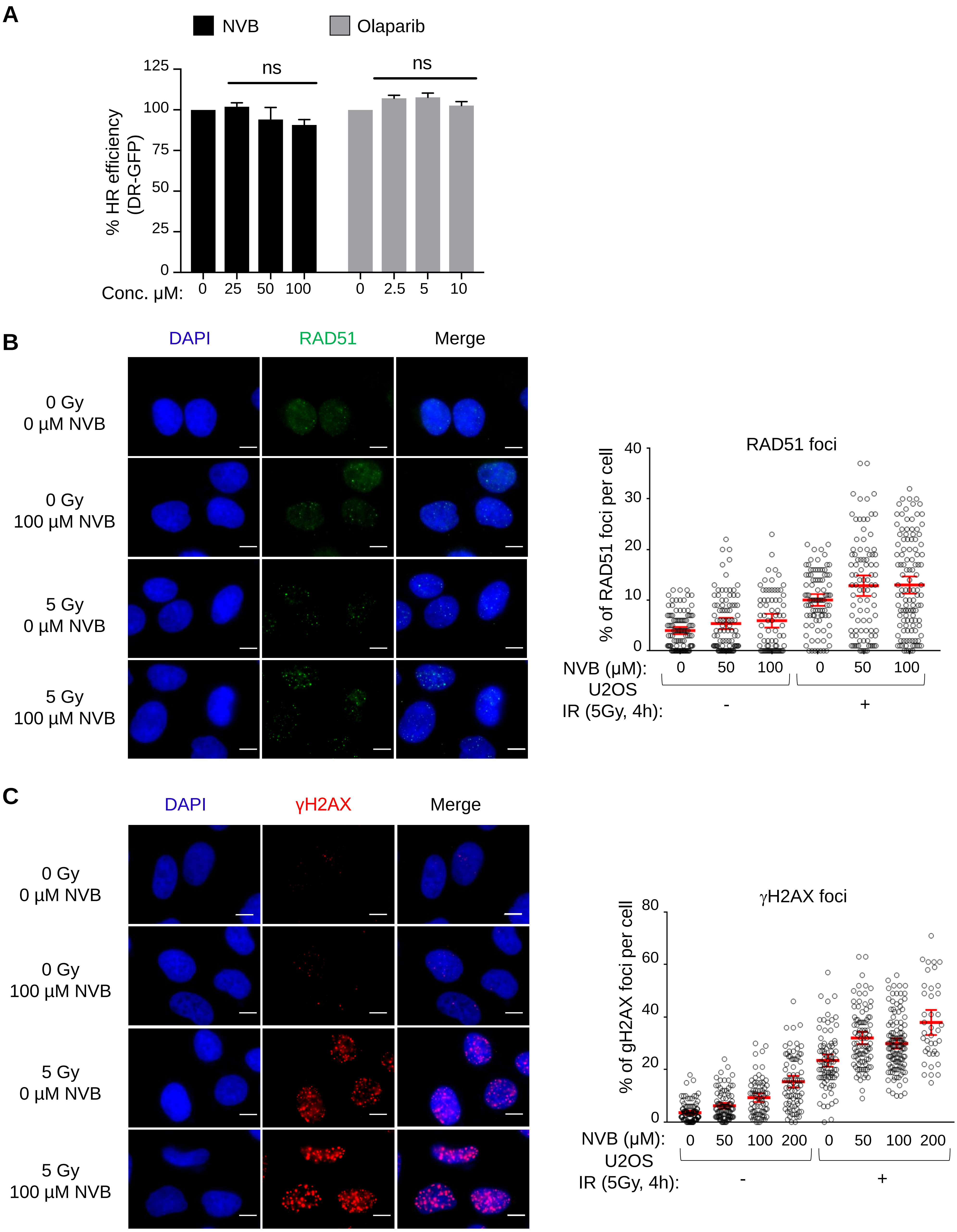

A
NVB
Olaparib
% HR efficiency
(DR-GFP)
125
100
75
50
25
0
ns
ns
Conc. µM:
0
25
50
100
0
2.5
5
10
B
DAPI
RAD51
Merge
0 Gy
0 µM NVB
0 Gy
100 µM NVB
5 Gy
0 µM NVB
5 Gy
100 µM NVB
RAD51 foci
% of RAD51 foci per cell
40
30
20
10
0
NVB (µM):
0
50
100
0
50
100
U2OS
IR (5Gy, 4h):
-
+
C
DAPI
γH2AX
Merge
0 Gy
0 µM NVB
0 Gy
100 µM NVB
5 Gy
0 µM NVB
5 Gy
100 µM NVB
γH2AX foci
% of gH2AX foci per cell
80
60
40
20
0
NVB (µM):
0
50
100
200
0
50
100
200
U2OS
IR (5Gy, 4h):
-
+

#### Fig S3

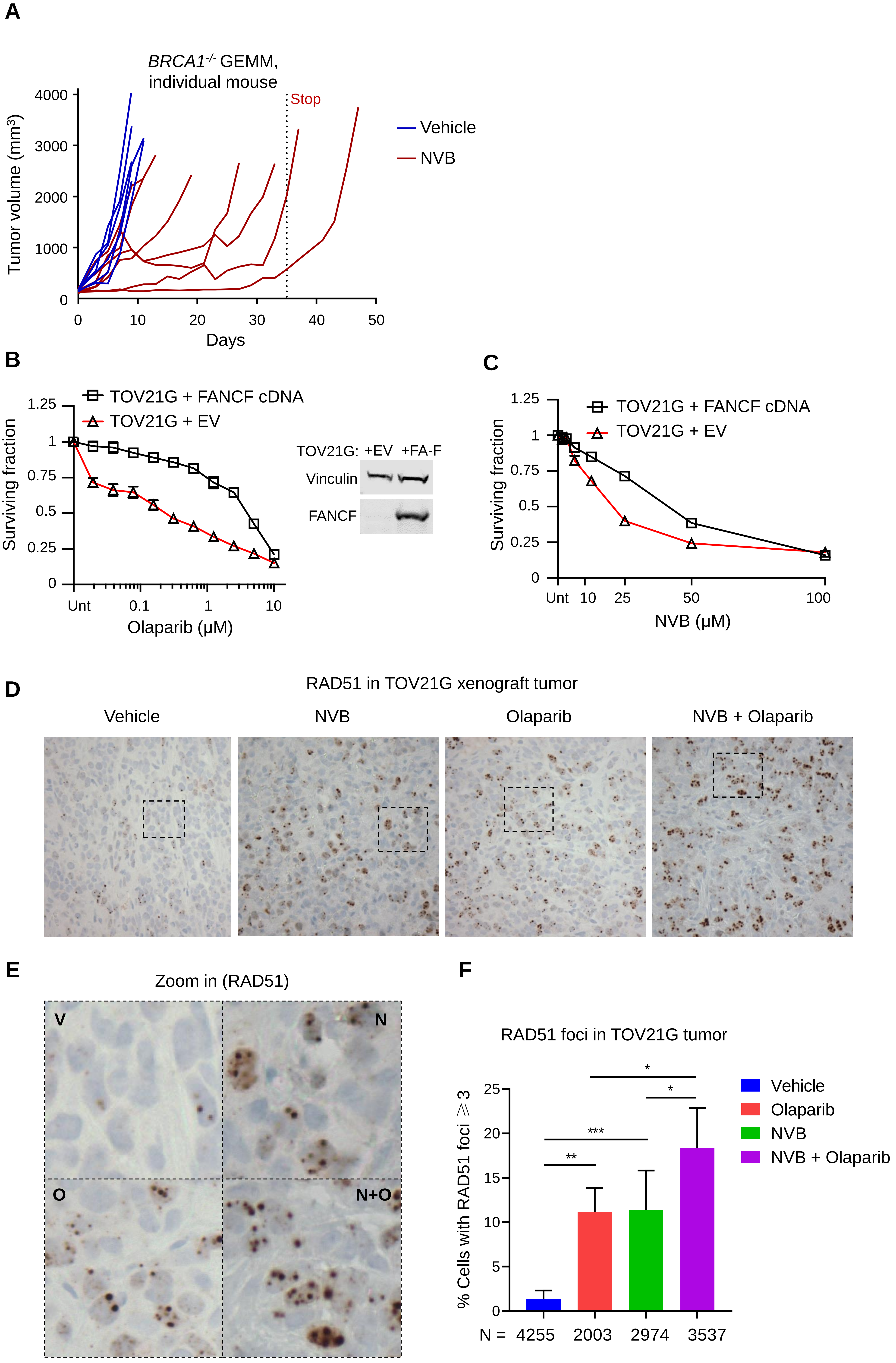
A
BRCA1-/- GEMM, individual mouse
Tumor volume (mm³)
Days
Stop
Vehicle
NVB
B
TOV21G + FANCF cDNA
TOV21G + EV
Surviving fraction
Olaparib (μM)
TOV21G: +EV +FA-F
Vinculin
FANCF
C
TOV21G + FANCF cDNA
TOV21G + EV
Surviving fraction
NVB (μM)
D
RAD51 in TOV21G xenograft tumor
Vehicle
NVB
Olaparib
NVB + Olaparib
E
Zoom in (RAD51)
V
N
O
N+O
F
RAD51 foci in TOV21G tumor
% Cells with RAD51 foci ≥ 3
N = 4255 2003 2974 3537
Vehicle
Olaparib
NVB
NVB + Olaparib

#### Fig S4

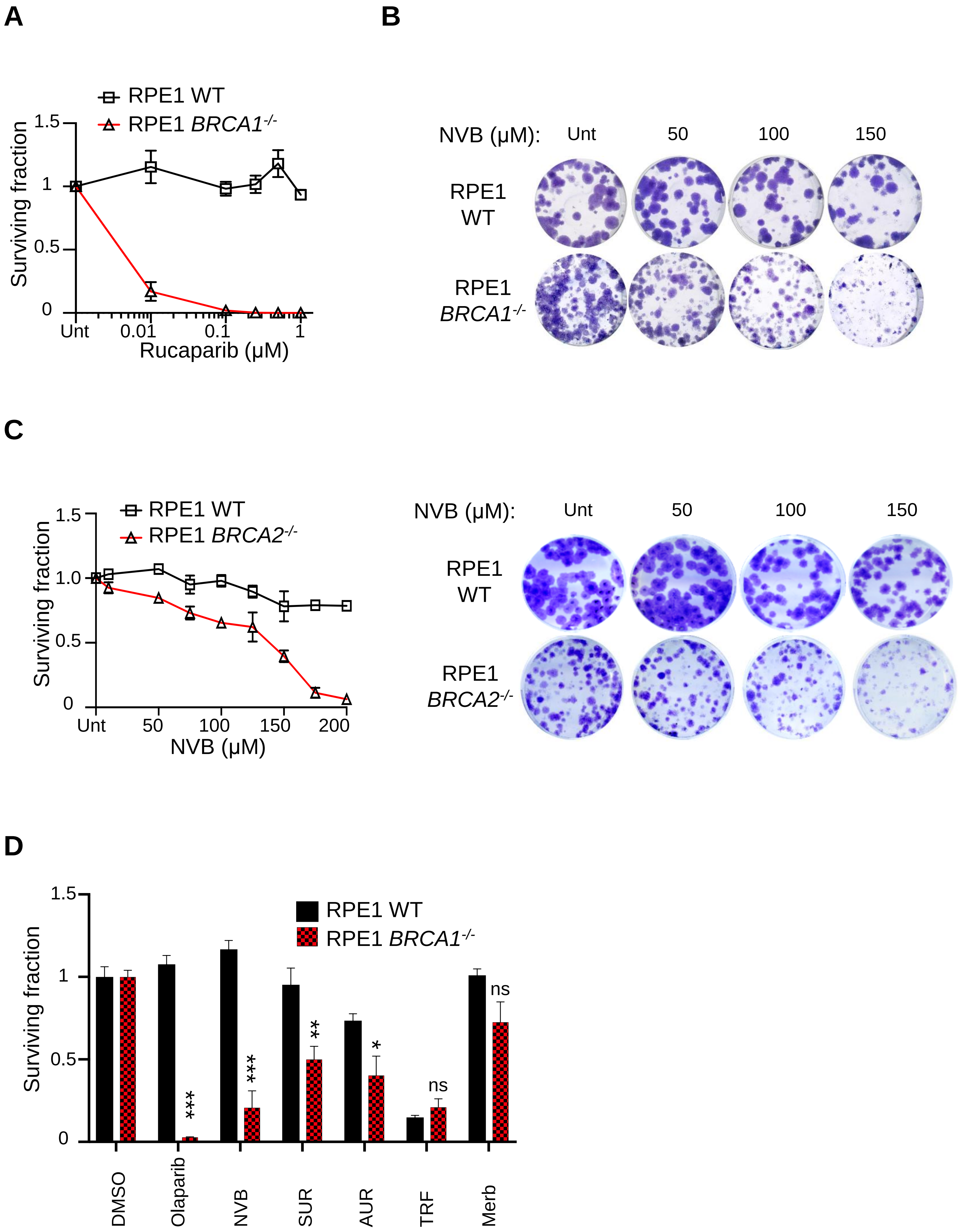
A
RPE1 WT
RPE1 BRCA1-/-
Surviving fraction
1.5
1
0.5
0
Unt
0.01
0.1
1
Rucaparib (µM)
B
NVB (µM):
Unt
50
100
150
RPE1 WT
RPE1 BRCA1-/-
C
RPE1 WT
RPE1 BRCA2-/-
Surviving fraction
1.5
1.0
0.5
0
Unt
50
100
150
200
NVB (µM)
NVB (µM):
Unt
50
100
150
RPE1 WT
RPE1 BRCA2-/-
D
RPE1 WT
RPE1 BRCA1-/-
Surviving fraction
1.5
1
0.5
0
***
***
**
*
ns
ns
DMSO
Olaparib
NVB
SUR
AUR
TRF
Merb

**Fig S5**

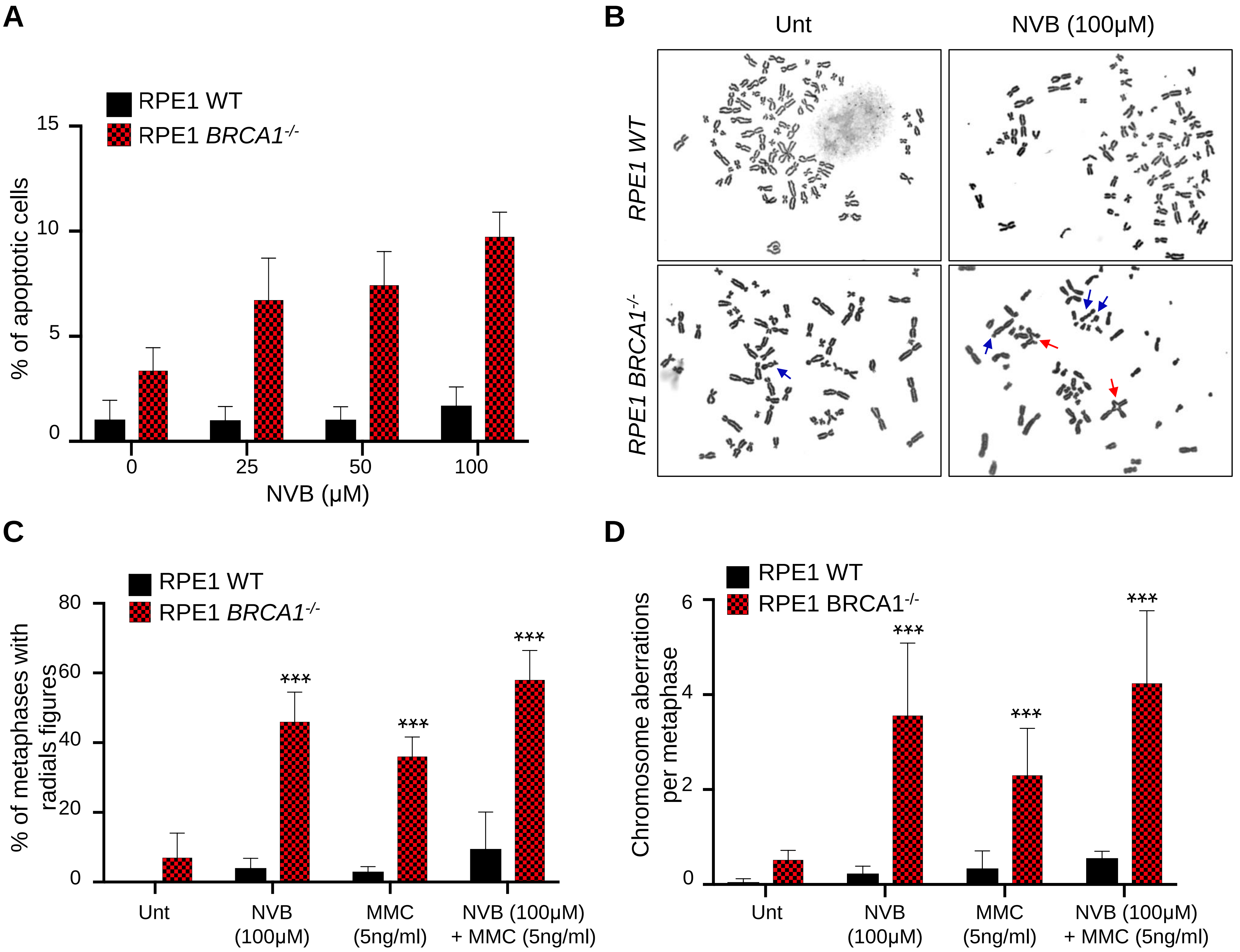

A
RPE1 WT
RPE1 BRCA1-/-
% of apoptotic cells
15
10
5
0
0
25
50
100
NVB (µM)
B
Unt
NVB (100µM)
RPE1 WT
RPE1 BRCA1-/-
C
RPE1 WT
RPE1 BRCA1-/-
% of metaphases with radials figures
80
60
40
20
0
***
***
***
Unt
NVB (100µM)
MMC (5ng/ml)
NVB (100µM) + MMC (5ng/ml)
D
RPE1 WT
RPE1 BRCA1-/-
Chromosome aberrations per metaphase
6
4
2
0
***
***
***
Unt
NVB (100µM)
MMC (5ng/ml)
NVB (100µM) + MMC (5ng/ml)

#### Fig S6

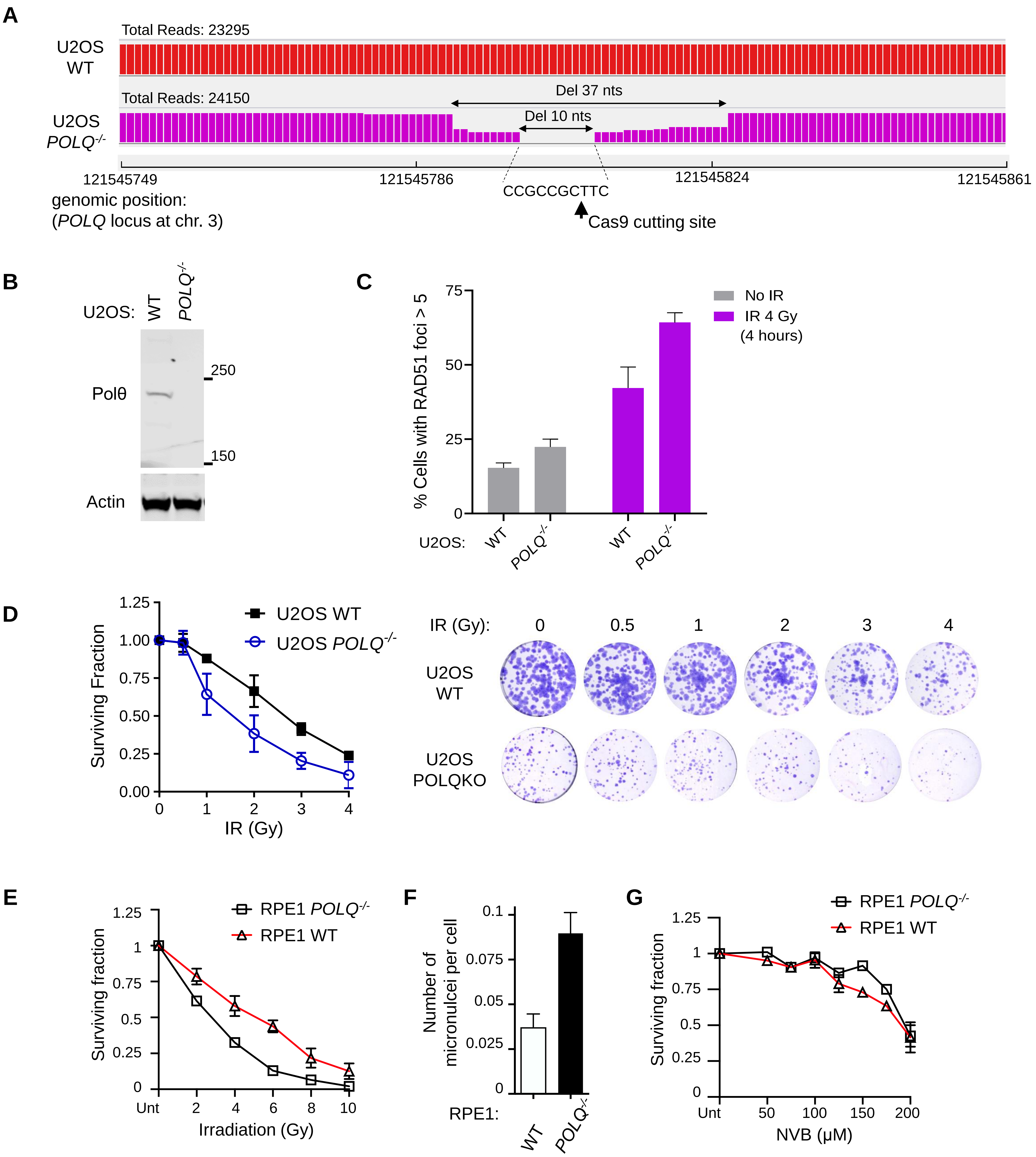
A
Total Reads: 23295
U2OS WT
Total Reads: 24150
U2OS POLQ-/-
Del 37 nts
Del 10 nts
121545749
121545786
121545824
121545861
CCGCCGCTTC
Cas9 cutting site
genomic position:
(POLQ locus at chr. 3)
B
U2OS: WT POLQ-/-
Polθ
250
150
Actin
C
% Cells with RAD51 foci > 5
75
50
25
0
No IR
IR 4 Gy
(4 hours)
U2OS: WT POLQ-/- WT POLQ-/-
D
Surviving Fraction
1.25
1.00
0.75
0.50
0.25
0.00
U2OS WT
U2OS POLQ-/-
IR (Gy)
IR (Gy): 0 0.5 1 2 3 4
U2OS WT
U2OS POLQKO
E
Surviving fraction
RPE1 POLQ-/-
RPE1 WT
Unt 2 4 6 8 10
Irradiation (Gy)
F
Number of micronulcei per cell
0.1
0.075
0.05
0.025
0
RPE1: WT POLQ-/-
G
Surviving fraction
RPE1 POLQ-/-
RPE1 WT
Unt 50 100 150 200
NVB (μM)

**Fig S7**

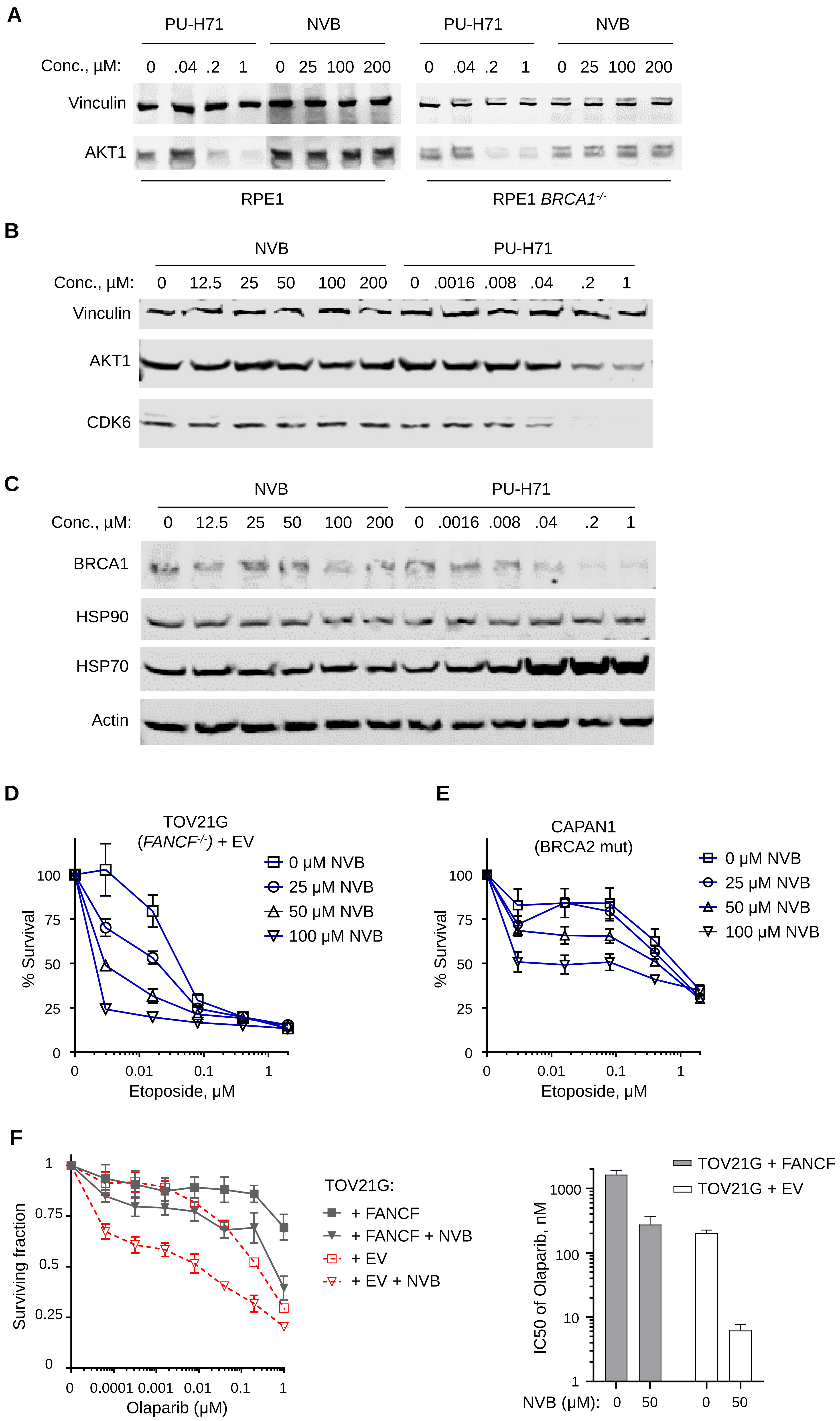
A
PU-H71
NVB
PU-H71
NVB
Conc., µM: 0 .04 .2 1 0 25 100 200 0 .04 .2 1 0 25 100 200
Vinculin
AKT1
RPE1
RPE1 BRCA1-/-
B
NVB
PU-H71
Conc., µM: 0 12.5 25 50 100 200 0 .0016 .008 .04 .2 1
Vinculin
AKT1
CDK6
C
NVB
PU-H71
Conc., µM: 0 12.5 25 50 100 200 0 .0016 .008 .04 .2 1
BRCA1
HSP90
HSP70
Actin
D
TOV21G
(FANCF-/-) + EV
0 µM NVB
25 µM NVB
50 µM NVB
100 µM NVB
% Survival
Etoposide, µM
E
CAPAN1
(BRCA2 mut)
0 µM NVB
25 µM NVB
50 µM NVB
100 µM NVB
% Survival
Etoposide, µM
F
Surviving fraction
Olaparib (µM)
TOV21G:
+ FANCF
+ FANCF + NVB
+ EV
+ EV + NVB
TOV21G + FANCF
TOV21G + EV
IC50 of Olaparib, nM
NVB (µM): 0 50 0 50

**Figure S8**

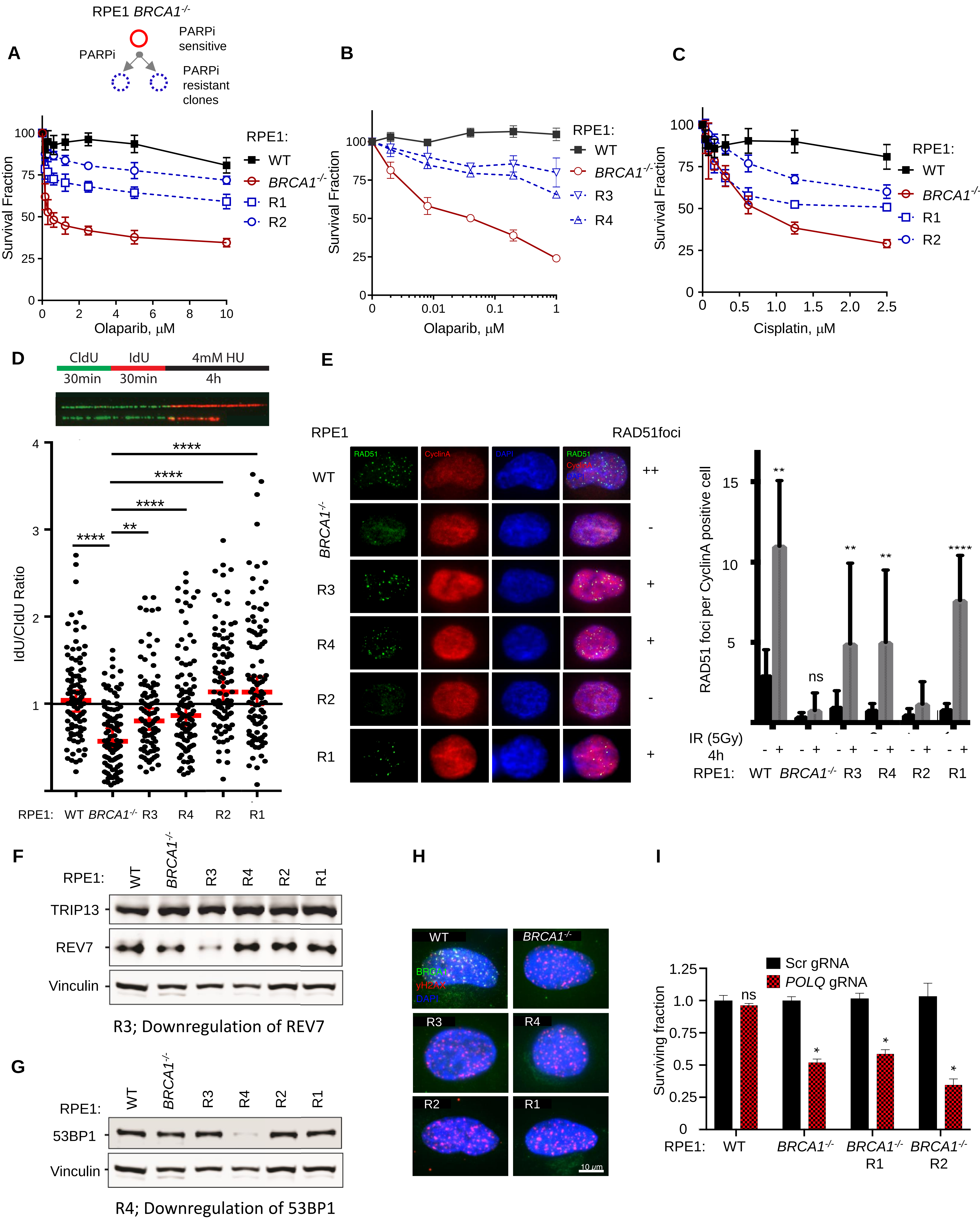

A
RPE1 BRCA1-/-
PARPi
PARPi sensitive
PARPi resistant clones
Survival Fraction
Olaparib, μM
RPE1:
WT
BRCA1-/-
R1
R2
B
Survival Fraction
Olaparib, μM
RPE1:
WT
BRCA1-/-
R3
R4
C
Survival Fraction
Cisplatin, μM
RPE1:
WT
BRCA1-/-
R1
R2
D
CldU
IdU
4mM HU
30min
30min
4h
IdU/CldU Ratio
RPE1: WT BRCA1-/- R3 R4 R2 R1
E
RPE1
RAD51foci
RAD51
CyclinA
DAPI
WT
BRCA1-/-
R3
R4
R2
R1
++
-
+
+
-
+
RAD51 foci per CyclinA positive cell
ns
IR (5Gy) 4h
RPE1: WT BRCA1-/- R3 R4 R2 R1
F
RPE1: WT BRCA1-/- R3 R4 R2 R1
TRIP13
REV7
Vinculin
R3; Downregulation of REV7
G
RPE1: WT BRCA1-/- R3 R4 R2 R1
53BP1
Vinculin
R4; Downregulation of 53BP1
H
WT
BRCA1-/-
BRCA1
γH2AX
DAPI
R3
R4
R2
R1
10 μm
I
Scr gRNA
POLQ gRNA
Surviving fraction
ns
RPE1: WT BRCA1-/- BRCA1-/- R1 BRCA1-/- R2

#### Fig S9

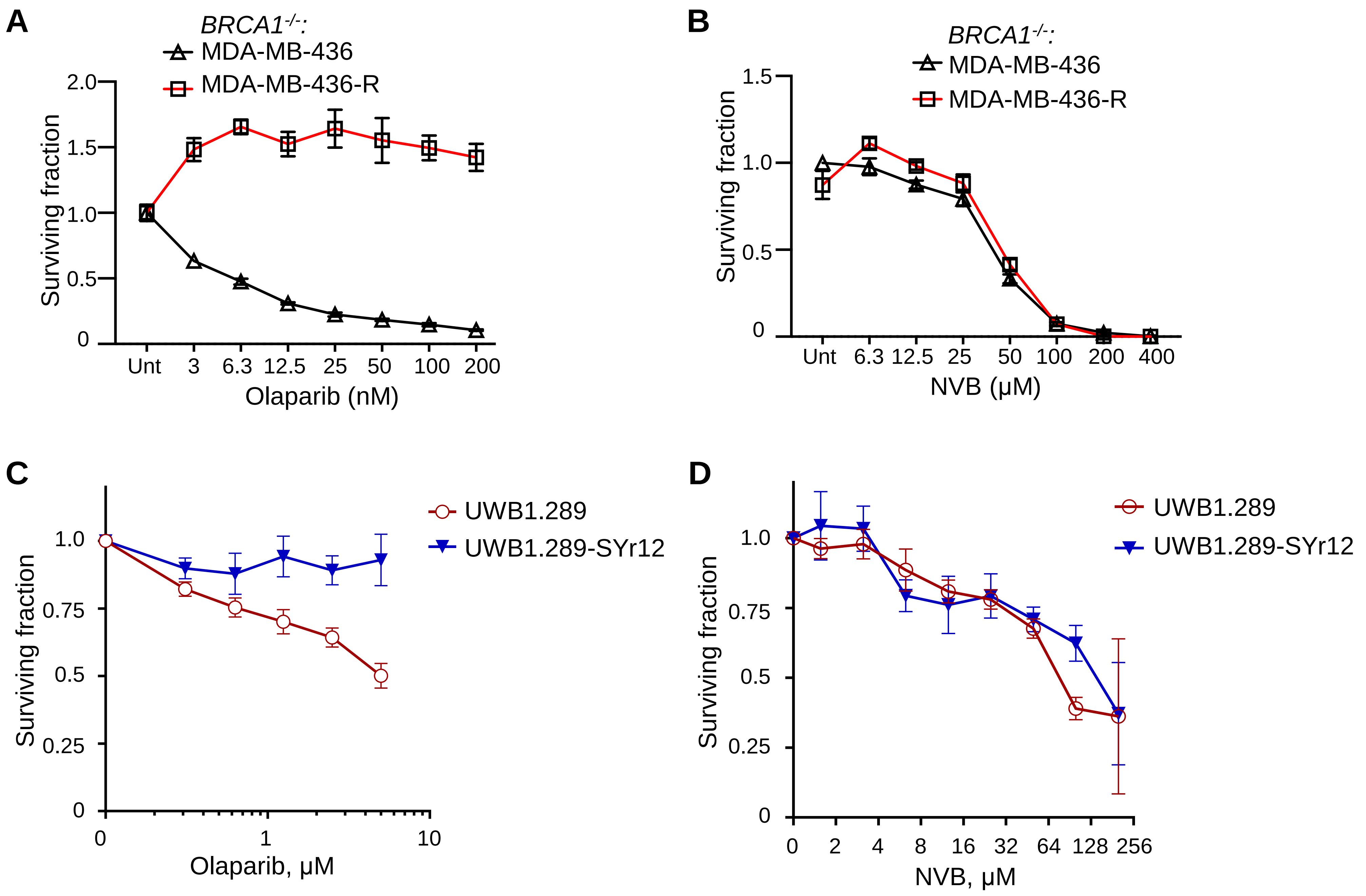
A
BRCA1-/-:
MDA-MB-436
MDA-MB-436-R
Surviving fraction
2.0
1.5
1.0
0.5
0
Unt
3
6.3
12.5
25
50
100
200
Olaparib (nM)
B
BRCA1-/-:
MDA-MB-436
MDA-MB-436-R
Surviving fraction
1.5
1.0
0.5
0
Unt
6.3
12.5
25
50
100
200
400
NVB (μM)
C
UWB1.289
UWB1.289-SYr12
Surviving fraction
1.0
0.75
0.5
0.25
0
0
1
10
Olaparib, μM
D
UWB1.289
UWB1.289-SYr12
Surviving fraction
1.0
0.75
0.5
0.25
0
0
2
4
8
16
32
64
128
256
NVB, μM

### Figure S10

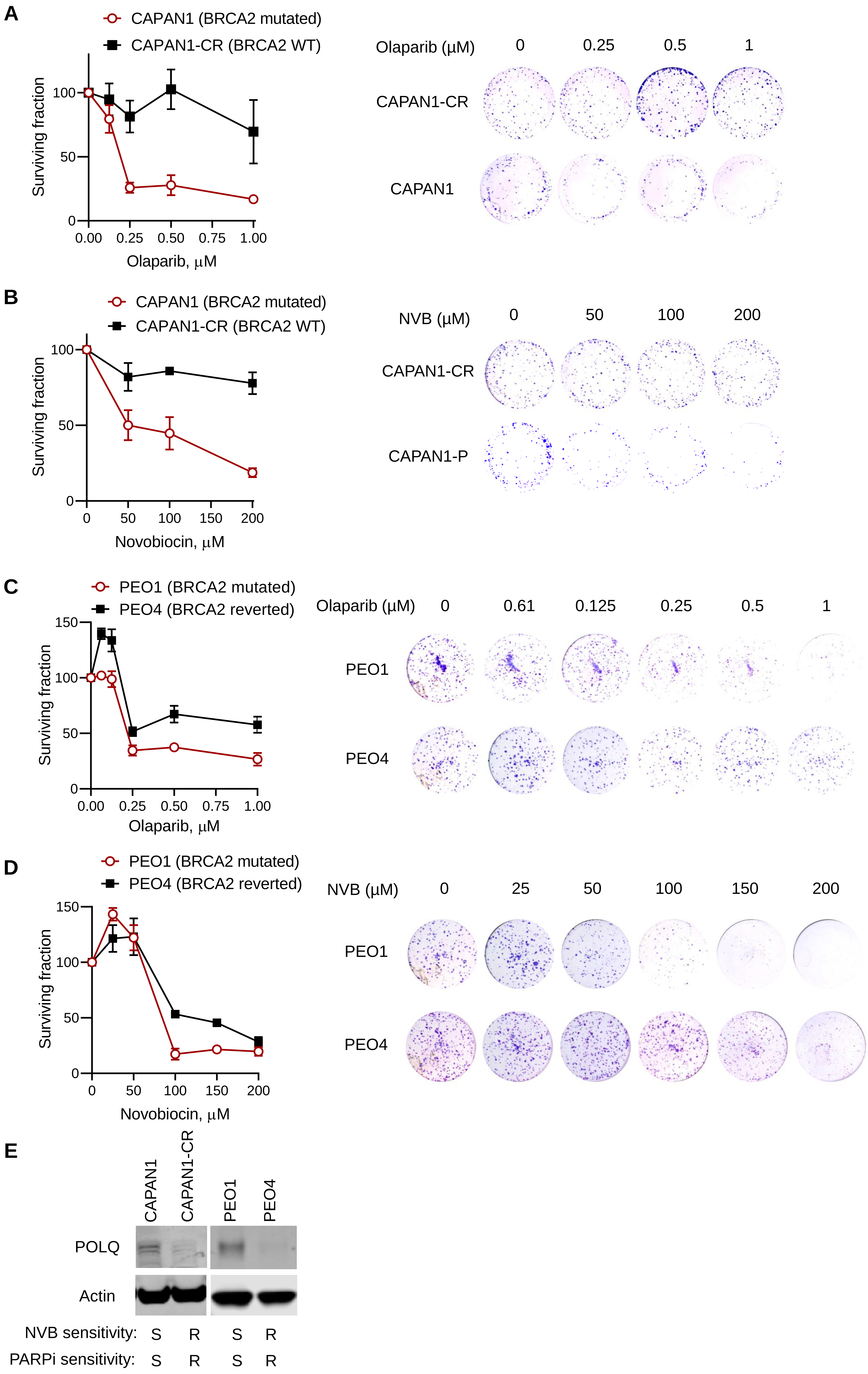
A
CAPAN1 (BRCA2 mutated)
CAPAN1-CR (BRCA2 WT)
Surviving fraction
100
50
0
0.00
0.25
0.50
0.75
1.00
Olaparib, µM
Olaparib (µM)
0
0.25
0.5
1
CAPAN1-CR
CAPAN1
B
CAPAN1 (BRCA2 mutated)
CAPAN1-CR (BRCA2 WT)
Surviving fraction
100
50
0
0
50
100
150
200
Novobiocin, µM
NVB (µM)
0
50
100
200
CAPAN1-CR
CAPAN1-P
C
PEO1 (BRCA2 mutated)
PEO4 (BRCA2 reverted)
Surviving fraction
150
100
50
0
0.00
0.25
0.50
0.75
1.00
Olaparib, µM
Olaparib (µM)
0
0.61
0.125
0.25
0.5
1
PEO1
PEO4
D
PEO1 (BRCA2 mutated)
PEO4 (BRCA2 reverted)
Surviving fraction
150
100
50
0
0
50
100
150
200
Novobiocin, µM
NVB (µM)
0
25
50
100
150
200
PEO1
PEO4
E
CAPAN1
CAPAN1-CR
PEO1
PEO4
POLQ
Actin
NVB sensitivity: S R S R
PARPi sensitivity: S R S R

#### SUPPLEMENTAL FIGURE LEGENDS

**Figure S1. Characterization of POLθ inhibitors obtained from the small-molecule screen, with *in vitro* biochemical assays and cell-based assays. A**, ADP-Glo assay for quantification of POLθ ATPase activity in various experimental conditions. **B**, A secondary screen demonstrated the potency of 60 of the 72 initial hits in reducing POLθ ATPase activity (below 70 %, *z*-score < -4). The screen was done in the presence and absence of ssDNA. The four most promising hits advancing for further analysis are labeled. Data shown are mean ± s.e.m (n=2). **C**, Titration of top putative ssDNA-binding inhibitors from the screen in reducing POLθ ATPase activity in the luminescent ADP-Glo assay. **D**, Titration of top putative POLθ-binding inhibitors from the screen in reducing POLθ ATPase activity in the luminescent ADP-Glo assay. **E**, A representative image from $^{32}P$ based radiometric POLθ ATPase assay with indicated small-molecule inhibitors from the screen. MTX, Mitoxantrone; SUR, Suramin; NNB, Novobiocin; AUR, Aurintricarboxylic Acid. The negative control is an inert small-molecule (Vandetanib) from the screen. **F-G**, Quantification of POLθ, SMARCAL1 and CHD1 ATPase activity in the presence of the indicated small-molecules. Data shown are mean ± s.d. (n=3). Statistics were performed using *t*-test with Welch's correction. ***$p < 0.001$, **$p < 0.01$, *$p < 0.05$. **H**, Conjugation of NVB to epoxy-activated Sepharose-6B (Sigma). NVB conjugated beads showed light yellow color. A structure of NVB is shown. **I**, Pulldown experiments with NVB-conjugated beads and cell lysate from HEK293T cells expressing GFP-tagged full-length POLθ. **J**, Competition assay between NVB-conjugated beads and free NVB when incubated with GFP-tagged full-length POLθ extracted from HEK293T cells. **K**, Thermal shift assay using HEK293T cell lysate incubated with Olaparib at the indicated temperatures.

**Figure S2. Novobiocin phenocopies POLθ depletion in human cells. A**, DR-GFP repair reporter assay (measures HR efficiency) in U2OS cells treated with indicated inhibitors. HR efficiency measured by GFP-positive cells was normalized to untreated samples. Data shown are mean ± s.e.m (n=3). **B**, Images and quantification of RAD51 foci in U2OS cells under increasing concentrations of NVB with or without gamma-irradiation (IR). **C**, Images and quantification of γH2AX foci in U2OS cells under increasing concentrations of NVB with or without IR. Data in (**B**) and (**C**) were pooled from two independent experiments. Each dot represents foci numbers in one cell.

**Figure S3. Efficacy of NVB in *BRCA1$^{-/-}$* GEMM and RAD51 PD study in TOV21G xenograft models. A.** NVB efficacy in the GEMM model (*Tp53$^{-/-}$Brca1$^{-/-}$* TNBC). Response of each tumor in each individual mouse is shown. Tumor chunks from GEMM mice were implanted in syngeneic FVB/129P2 mice, which were treated with PBS or 100 mg/kg NVB twice a day via IP for 5 weeks. **B**, Olaparib sensitivity of TOV21G (+ EV) or FANCF-complemented TOV21G (+ FANCF cDNA). **C,** Novobiocin sensitivity of TOV21G (+ EV) or FANCF-complemented TOV21G (+ FANCF cDNA). **D-F**. Immunohistochemical (IHC) study of the pharmacodynamic biomarker RAD51 after NVB and/or olaparib treatment in TOV21G tumors. **D**. Representative images of RAD51 IHC staining in xenografted TOV21G tumor cells. Tumor bearing NU(NCr)-Foxn1nu mice were treated with indicated drugs for 18 days before tumors were taken. FFPE tissue sections of the tumors were stained using an anti-RAD51 antibody and representative images (40X) are shown. **E**. Zoom-in of the IHC images in **(D)** to show RAD51 foci in detail. **F**. Quantification of RAD51 foci positive cells in TOV21G tumors. Cells with three or more RAD51 foci were counted as positive cells. Three tumor samples from each group were processed and analyzed, and the total number of cells counted were shown (N). Statistical analysis was performed using one-way ANOVA in Prism, *, $p < 0.05$; **, $p < 0.01$; ***, $p < 0.001$.

**Figure S4. Effects of novobiocin in *BRCA1$^{-/-}$* and *BRCA2$^{-/-}$* cells. A**, Clonogenic survival of *BRCA1$^{-/-}$* and WT RPE1 cells under increasing concentrations of the PARPi rucaparib. The survival fraction is normalized to the untreated sample (Unt). **B**, Representative images from clonogenic survival of *BRCA1$^{-/-}$* and WT RPE1 cells under increasing concentrations of the POLθ inhibitor NVB. Three independent experiments were performed. **C**, Clonogenic survival of *BRCA2$^{-/-}$* and WT RPE1 cells under increasing concentrations of novobiocin. Quantifications and representative images and were shown. The survival fraction is normalized to the Unt samples. **D**, Cell viability assay (CellTiter-Glo) in *BRCA1$^{-/-}$* and WT RPE1 cells treated with indicated POLθ inhibitors (50 μM) or with the PARPi Olaparib (50 nM) as control. Cells were treated twice on days 1 and 4 and cell viability was measured on day 7. Data were mean ± SD, n = 3. Statistics analyses were *t*-test.

**Figure S5. NVB induces apoptosis and chromosome aberrations in *RPE1 BRCA1$^{-/-}$*. A**, Quantification of apoptosis in *BRCA1$^{-/-}$* and WT RPE1 cells under increasing concentrations of NVB. Mean ± s.d., n = 3. **B,** Representative images of chromosomal aberrations (blue arrows) including radial figures (red arrows) in RPE1 (WT or *BRCA1*$^{-/-}$) cells treated with NVB. **C-D**, Quantification of radial figures (**C**) and total chromosome aberration (**D**) in *BRCA1*$^{-/-}$ and WT RPE1 cells treated with NVB alone or in combination with mitomycin C (MMC). Data in **C-D** are mean ± s.d. of n = 3 independent experiments. Statistical significance was determined by one-way ANOVA multiple comparisons using uncorrected Fisher's LSD test. ***, $p < 0.001$.

**Figure S6. Verification of *POLQ*$^{-/-}$ cells. A-D**, Verification of the U2OS-*POLQ*$^{-/-}$ cell line. **A**, mapping the genetic alteration (deletions) in the U2OS *POLQ* knockout clone generated by CRISPR-Cas9. The targeted region was sequenced by Next Gen sequencing (number of reads are shown), and the deletions were mapped to *POLQ* locus of human genome on chromosome 3. **B**, A Western blot showing POLθ protein expression in WT and *POLQ*$^{-/-}$ U2OS cells. **C**, RAD51 foci assay in WT or *POLQ*$^{-/-}$ U2OS cells before and after (4 h) of 4 Gy IR. **D**, IR sensitivity of WT and *POLQ*$^{-/-}$ U2OS cells in clonogenic assays. **E-F**, Verification of the RPE1-*POLQ*$^{-/-}$ cell line. **E**, IR sensitivity of WT and *POLQ*$^{-/-}$ RPE1 cells in clonogenic assays. **F**, Quantification micronuclei, a hallmark of POLQ loss, in WT and *POLQ*$^{-/-}$ RPE1 cells. **G**, *POLQ*$^{-/-}$ RPE1 cells show resistance to NVB when compared to WT.

**Figure S7. NVB inhibits POLθ but not HSP90 or TOP2 in human cells. A**, HSP90 client degradation assay in RPE1 and RPE1-*BRCA1*$^{-/-}$ cells. Cells were treated with a potent HSP90 inhibitor PU-H71 or NVB for 48 hours, and then cells were collected for Western blot analysis of the HSP90 client AKT1. **B-C**, HSP90 client degradation assay in MCF7 cells. Cells were treated with the potent HSP90 inhibitor PU-H71 or NVB for 24 hours, and then cells were collected for Western blot analysis of the HSP90 client proteins AKT1, CDK6 (**B**) and BRCA1 (**C**). Levels of HSP90 and HSP70 were also analyzed after NVB or PU-H71 treatment (**C**). **D-E**, Combination effect of etoposide and novobiocin in killing TOV21G cells (**D**) and CAPAN1 cells (**E**), showing their non-epistatic effects. **F**, CellTiter-Glo cell viability assay of empty vector (EV) and FANCF-complemented TOV21G cells, in the presence of olaparib, novobiocin or both. $IC_{50}$ values of olaparib in TOV21G cells with or without NVB are shown on the right.

**Figure S8. Characterization of PARP inhibitor resistant clones derived from RPE1-*BRCA1*$^{-/-}$. A-C**, Olaparib (**A-B**) and cisplatin (**C**) sensitivity of WT and *BRCA1*$^{-/-}$ RPE1 cells and the PARPi resistant clones of *BRCA1*$^{-/-}$ RPE1. Data shown are mean ± SD from n ≥ 3 of CellTiter-Glo assay measurements. **D**. DNA fiber assay to measure the replication fork stability of the PARPi resistant clones. **E**. RAD51 foci analysis in WT and *BRCA1-/-* RPE cells and the PARPi resistant clones after irradiation (5Gy), to determine restoration of RAD51 foci in R clones. RAD51 was stained 4 hours after IR. **F**. A Western blot of WT and *BRCA1-/-* RPE cells and the PARPi resistant clones, using TRIP13 and REV7 antibodies. **G**. A Western blot of WT and BRCA1-/- RPE cells and the PARPi resistant clones, using an antibody against 53BP1 (an essential NHEJ protein). **H**. Immunofluorescence staining of BRCA1 in WT and *BRCA1-/-* RPE cells and the PARPi resistant clones after irradiation. No BRCA1 foci was observed except in WT RPE1 cells. **I**, CellTiter-Glo assay to determine survival of PARPi-resistant and parental *BRCA1*$^{-/-}$ RPE1 cells upon CRISPR knockout of the *POLQ* gene. Data are mean ± s.e.m., with n = 4. Statistical analysis was *t*-test, *, $p < 0.05$; ns, not significant.

**Figure S9. PARPi resistant MDA-MB-436 and UWB1.289 cells are sensitive to NVB, and NVB has synergy with PARPi in TOV21G cells. A-B**, CellTiter-Glo cell viability assay in parental MDA-MB-436 (*BRCA1*-mutated) cells and a PARPi-resistant subclone (MDA-MB-436-R) under increasing concentration of Olaparib **(A)** or NVB **(B)**. Mean ± s.e.m. are shown, with n = 6. **C-D**, Olaparib (**C**) and NVB (**D**) sensitivity of UWB1.289 and a PARPi resistant UWB1.289 clone (UWB1.289-YSR12) in CellTiter-Glo assays. Data shown are mean ± SD, n = 4.

**Figure S10. Cancer cells with BRCA2 mutations reverted to WT are not sensitive to novobiocin. A**. Olaparib sensitivity of CAPAN1 (BRCA2 mutated) and CAPAN1-CR cells (BRCA2 CRISPR edited back to wild type). **B**. NVB sensitivity of CAPAN1 and CAPAN1-CR cells. **C.** Olaparib sensitivity of PEO1 (BRCA2 mutated) and PEO4 cells (BRCA2 restored). **D**. NVB sensitivity of PEO1 and PEO4 cells. Experiments in **A-D** were clonogenic survival assays. Data shown are Mean ± S.D., n = 2 (**A-B**) and n= 3 (**C-D**). **E**. A Western blot shows POLQ expression in BRCA2-decient cells lines (PEO1 and CAPAN1) and their counterparts with BRCA2 reverted to wild type (PEO4 and CAPAN1-CR). Corresponding NVB and olaparib sensitivities of each cell line were labeled.

#### MATERIALS AND METHODS

##### Compounds, inhibitors and antibodies

Chemical compounds, including PARP inhibitors Olaparib and Rucaparib, Novobiocin, etoposide, and PUH71 were purchased from Selleckchem. Chemicals were dissolved in DMSO and kept in small aliquots at -80°C. The key sources of antibodies are: POLθ (Sigma), γH2AX (Millipore), RAD51 (Santa Cruz Biotechnology, SCB), CDK6 (SCB), AKT1 (SCB), HSP70 (SCB), HSP90 (SCB), and FANCF (Everest Biotech).

##### Protein purification

A POLθ fragment (ΔPol) containing the ATPase domain with a RAD51 binding site (amino acids 1 to 987) was cloned into pFastBac-C-Flag and purified from baculovirus-infected SF9 insect cells. The POLθ-pFastBac-C-Flag vector was transformed into DH10bac E. coli, and bacteria were plated on an IPTG and X-Gal containing plates and recombinant bacmid DNA was purified using the Qiagen midiprep kit. POLθ bacmid was transfected into the SF9 cells grown in supplemented Grace’s Insect Medium using the Cellfectin II reagent. Baculovirus excreted from the cells and the surrounding media were collected after three days (P1). SF9 cells were grown to 90-100% confluence in 15 cm plates. 150 μl of P1 virus was added to the plate covered with 25 mL Grace’s Insect Medium. The surrounding media containing P2 virus was collected after 3 days. Then, SF9 cells were grown in 1 L spinner flasks at a volume of 500 mL and a density of 1.5-2 million cells/mL. At the concentration of 1.5-2 million cells/mL, 20 mL of P2 virus was added to the 500ml culture. In spinner flask culture Grace’s media were supplemented with Pluronic F68 (Thermo Fisher) to limit shearing. Three days after infection, cells were spun down and frozen with liquid nitrogen. Cells were lysed in 500 mM NaCl lysis buffer (500 mM NaCl, 0.01 % NP40, 0.2 mM EDTA, 20% Glycerol, 1 mM DTT, 0.2 mM PMSF, 20 mM Tris [pH 7.6]) supplemented with Halt protease inhibitor cocktail (Thermo Fisher) and Calpain I inhibitor (Roche). The cell lysates were incubated with M2

Flag beads (Sigma) for 3 hours at 4 $^{o}$C on a spinning wheel. The M2 Flag beads were washed 3 times lysis buffer and protein was then eluted in lysis buffer supplemented with 0.2 mg/ml of Flag peptide (Sigma). The protein was then concentrated in lysis buffer using 10 kDa centrifugal filters (Amicon), quantified and flash-frozen in small aliquots in liquid nitrogen and stored at -80°C.

**ADP-Glo ATPase assay**

ATPase activity of a POLθ (ΔPol) was measured using the ADP-Glo kinase assay (Promega). A 10 µl mix containing 10 nM of POLθ-ΔPol protein, 600 nM of 30mer single-stranded DNA substrate, 40 mM Tris-HCl buffer (pH 7.6), 20 mM MgCl2, 0.1 mg/ml BSA, and 1 mM DTT was added to each reaction well in a black 384 well plate (Corning). Then, 0.1 µL of a chemical inhibitor or negative control (DMSO) was added to each well. 3 µl of 433.33 µM purified ATP (from ADP-Glo kit) or reagent buffer (No ATP positive control) was then added to each reaction well. DMSO wells represented 0% inhibition while no-ATP wells represented 100% inhibition. Plates were covered with an aluminum seal, stacked, and incubated at room temperature overnight (~16 hours). The plates were stored in a humid, plastic box to decrease evaporation. The following day, 6.5 µl of ADP-Glo reagent (from kit) was added to each reaction well to remove unhydrolyzed ATP. After 1-hour incubation, 13 µl of Kinase detection reagent was then added to the wells, and plates were incubated for another hour. Finally, ATP hydrolysis was quantified by luminescence measured on a plate reader.

**High Throughput Screening**

The ATPase activity based high throughput small-molecule screen was performed at ICCB Longwood Screen Facility at Harvard Medical School. On the day of screening, Combi machine was used to dispense 10 µl aliquots of a master mix containing 600 nM of 30-mer single-stranded DNA substrate and 10 nM of POLθ protein (ΔPol) into each well of a 384 well plate (Corning 3820). Small-molecules were added to wells in the 384 well plates by Seiko Pin transfer robot. Plates were covered with aluminum seals and spun down at 1000 rpm and sat for 1 hour to allow small-molecules to bind protein. 3 μL of 433.33 μM ultra-pure ATP was added to every well except no-substrate control wells. Plates were stored overnight (16 hours) in a humidified plastic chamber. Next day, 6.5 μL of ADP-Glo reagent (Promega) was added to every well using the Combi machine. After one-hour incubation, 13 μL of Kinase Detection Reagent (Qiagen) was added to every well and incubated for another hour. Data was collected using the Envision plate reader by measuring luminescence signal.

**Statistical Analysis of the high throughput screen**

The inhibition strength of screened compounds was evaluated using Z-score analysis. Since most compounds did not affect POLθ-ATPase activity, data for all compounds was used to calculate plate mean (μ) and standard deviation (σ). Every compound is then assigned a Z-score using the equation $z = (x – \mu) / \sigma$. Percentage inhibition analysis. Percentage inhibition of compounds was determined by normalizing to the DMSO control. Wells with DMSO were averaged to find the average signal without inhibitor, and a different DMSO average was calculated for each plate. The % activity was calculate using the following formula: % activity = [(signal with compound) / (average signal with DMSO)] x 100%.

**$^{32}$P-based ATPase activity assay**

POLθ protein (ΔPol, aa 1-987) was pre-incubated with inhibitors and ssDNA for 30 minutes (min) in reaction buffer (20 mM HEPES-KOH [pH 7.6], 100 mM KCl, 5 mM MgCl2, 0.25 μg/μl BSA, 0.05 mM EDTA, 0.5 mM DTT, 3% glycerol, and 0.01% NP-40). The substrate ATP (500 μM cold ATP and 1 μCi of [γ-$^{32}$P]-ATP as a tracer) was added to start the reactions. The total volume of the reactions was 20 μl. Aliquots (2 μl) of the reactions were removed and immediately dotted to a dry PEI-cellulose plate (Sigma) at indicated time points. The samples were resolved by thin-layer chromatography (TLC) in 4.5% formic acid supplied with 0.5 M lithium chloride. TLC plate was then air-dried and exposed to a phosphor screen for visualization of radioactivity by a Personal Molecular Imager (PMI) System (Bio-Rad). Quantification of the [$\gamma$-$^{32}$P]-ATP and the released inorganic phosphate ($^{32}$Pi) was performed using Quantity One software. After background subtraction, the fraction of hydrolyzed ATP was calculated.

**Thermal Stability Assays (TSA)**

HEK293T cells overexpressing eGFP-POLθ were lysed in IP buffer (Thermo Fisher, #87788) supplemented with PMSF, protease and phosphatase inhibitors. 50 μL of cell lysate was aliquoted to each tube and NVB or DMSO was added. The mixture was incubated for 15 min at room temperature, heated at indicated temperature (37 $^{o}$C to 55 $^{o}$C) for 3 min. The temperature was subsequently reduced to room temperature for another 5 min. Next, the mixture was centrifuged at 20,000 g for 15 minutes. The supernatant was analyzed by Western blot.

**Preparation of Novobiocin–Sepharose 6B Beads**

NVB–Sepharose 6B Beads were prepared similarly as previously described (*1*). One gram of epoxy-activated Sepharose 6B (Sigma-Aldrich E6754) was washed thoroughly and swollen in 50 mL of distilled water for 1 hour at room temperature. The resin was washed once with coupling buffer (0.3 M Sodium Carbonate [pH 9.5]). The washed resin was mixed with 100 mg of NVB in 10 mL of coupling buffer and incubated at 37 °C with rotation overnight. Next day, the resin was washed with coupling buffer three times to remove excess NVB. The remaining epoxy-active groups were blocked with 1 M ethanolamine (in coupling buffer) for 8 hours at 30 °C with gentle rotation. The beads were sequentially washed with coupling buffer, 0.5 M NaCl in coupling buffer, distilled water, 0.5 M NaCl in 0.1 M sodium acetate (pH 4), and twice in distilled water. The beads were re-suspended in 25 mM HEPES (pH 7.6), 200 mM KCl, 10% ethylene glycol, and 1 mM EDTA. Inactivated Novobiocin–Sepharose 6B Beads (NVB inact) were obtained by light and temperature exposure.

**GFP reporter-based DNA repair assays**

U2OS cells containing the DR-GFP (for HR) and EJ-2 (for MMEJ) repair substrates were gifts from Dr. Jeremy Stark at Beckman Research Institute of the City of Hope (*2*). To measure the repair efficiency, $1\times10^5$ cells were plated in each well of a 12-well plate. Indicated compounds or DMSO were added to the medium for 24 hours, and cells were infected with Adenoviruses expressing the I-SceI enzyme. Two days after infection, cells were trypsinized and GFP-positive cells were quantified by flow cytometry (Beckman).

**Cell culture and transfection**

Human cells were maintained in culture media (DME-HG/F-12 for WT and knockouts RPE1 cells; RMPI for MDA-MB-436; DMEM for U2OS) supplemented with 10% FBS and 1% Pen-Strep. Full length POLθ was cloned by Gibson assembly in pCAG-GFP (Plasmid #11150) and co-transfected with Super PiggyBac Transposase Expression Vector (#PB210PA-1, System Biosciences) using Lipofectamine LTX with Plus Reagent in RPE-P53.

**RAD51 and γH2AX focus formation assay**

$2.5 \times 10^5$ U2OS cells were seeded in a 35 mm MatTek dish in the morning of day 1. In the afternoon, NVB was added to the medium at indicated concentrations. 24 hours later, cells were treated with 5 Gy of irradiation. 4-8 hours after irradiation, cells were extracted and fixed with 1% PFA, 0.5% methanol, and 0.5% TritonX-100 in PBS for 20 min at room temperature on a shaker. Fixed cells were washed with PBS twice and blocked with BTG buffer (1 mg/mL BSA, 0.5% TritonX-100, and 3% of goat serum, 1 mM EDTA) for 1 hour. Cells were then incubated with a rabbit Rad51 antibody (Santa Cruz Technology sc-8349 or Cell Signaling Technology #8875) and mouse γH2AX antibody (Millipore) in BTG buffer. After wash with PBS three time, cells were incubated with Alexa-488 or Alexa-594 conjugated secondary antibodies (Life Technologies) in BTG buffer for 1 hour. The dishes were washed with BTG buffer once and PBS twice, and mounted with mounting solution with DAPI for immunofluorescence microscopy. Images of random fields were taken under a 63x oil lens.

**Clonogenic survival assay**

Cells were seeded, at 2 different concentrations, into each well of a 6-well plate (100 and 200 cells for WT RPE1; 500 and 1000 cells for HR-deficient cells). Next day, cells were treated with the indicated doses of inhibitors. For combination studies, cells were treated first with Novobiocin (100 µM) or DMSO for 24h, washout and then release in fresh media containing Novobiocin (100 µM) or DMSO with increasing concentration of PARPi. Cells were allowed to grow in drug-containing medium for 12 to 14 days. Cells were fixed and stained with 0.2% crystal violet in methanol for 30 min (Crystal Violet solution, Sigma HT90132), and rinsed with distilled water three times. The stained dishes were air-dried, and the number of colonies (>50 cells) was counted in each well.

**Gene knockouts**

BRCA1 and BRCA2 knockouts were generated in RPE-1 cells with TP53 knockout (RPE-P53). TP53 knockout was generated by co-transfection of Cas9 (pSpCas9(BB)-2A-GFP Addgene #48138) (PX458) and TP53 sgRNA vectors (gRNA cloning vector Plasmid Addgene #41824) by Lipofectamine LTX with Plus Reagent (Thermo Fisher). Single colonies were screened by Western blotting. RPE1-P53 BRCA2$^{-/-}$ cells were obtained by co-transfection of Cas9 and two sgRNA vectors targeting introns flanking exon 2 of the *BRCA2* gene. After subcloning into 96 wells plate, single colonies were screened by PCR. Briefly, DNA was extracted by adding 30µl of a 50mM NaOH solution per well after which the plate was put at 95°C for 10 minutes. Then, 2.5µl of 1M Tris-HCL PH 7.5 was added per well and 2µl of the extracted DNA was used to perform PCR screening (Terra PCR Direct

Polymerase, Takara Clontech). Clones were selected for the presence of deletion PCR band and/or absence of 5' and 3' junction PCR band. The SgRNA sequences used were Tp53: GGCAGCTACGGTTTCCGTC; BRCA2-2: GGTAAAACTCAGAAGCGC; BRCA2-3: GCAACACTGTGACGTACT. The PCR primer sequences used for genotyping were the following: (Deletion PCR: seqBRCA2-For: GCTGTATTCCGAAGACATGCTGATGG; seqBRCA2-Rev: TTGTTCCTACTGCTAGTCAAGGG), (5' Junction PCR: seqBRCA2-For + exon2-Rev: TACCTACGATATTCCTCCAATGCTT), (3' Junction PCR: seqBRCA2-Rev + exon2-For: AAGCATTGGAGGAATATCGTAGGTA). *POLθ* knockout cells were generated in U2OS cells. Cas9 was introduced into U2OS cells by lentiviral infection using the lentiCas9-Blast plasmid (Addgene #52962), and stable Cas9-expressing U2OS cells (U2OS-Cas9) were obtained after blasticidin selection. SgRNA oligos designed to target the genomic sequence GATTCGTTCTCGGGAAGCGG of *POLθ* (exon 1) were annealed and inserted into LentiGuide-Puro plasmid (a gift from Feng Zhang, Addgene #52963). After lentivirus infection, U2OS-Cas9 cells were selected by puromycin for 3 days, and the survived cells were trypsinized and seeded sparsely to form single-cell colonies. Colonies were picked and expanded, and genomic DNA from individual clones was isolated by QIAamp DNA Mini Kit (Qiagen). PCR was performed to amplify the gRNA-targeted region (216 bp), using the following primers: Forward, GGAGGACGCTGGGACTGTGGC; Reverse, CTGCAGCTGCGGCCTTCAGGC. PCR products were cleaned by PCR purification kit (Qiagen) and submitted for Next-Gen Sequencing (Amplicon sequencing provided by Genewiz). The sequencing reads were mapped to the human genome at *POLθ* locus and the results were visualized by Integrative Genomics Viewer (IGV, Broad Institute). Clones with homologous deletion of *POLθ* gene were confirmed by Western blot analysis.

**Laser micro-irradiation**

RPE1-P53 cells overexpressing eGFP-POLθ were plated in a 35 mm µ-Dish (Ibidi). The day after, DMSO, Rucaparib (1 µM) or Novobiocin (200 µM) were added to the cells for 24 hours. Micro-irradiations were performed with a two photons laser (800 nm) on an inverted Laser Scanning Confocal Microscope equipped with Spectral Detection and Multi-photon Laser (LSM880NLO/Mai Tai Laser - Zeiss/Spectra Physics) with Airyscan module. Images were then analyzed using Fiji and statistical analyses were performed using Prism7.

**Mouse Xenograft and PDX studies**

All animal experiments were conducted in accordance with Institutional Animal Care and Use Committee-approved protocols at Beth Israel Deaconess Medical Center and Dana-Farber Cancer Institute.

**TOV21G xenograft study:** Female NU(NCr)-*Foxn1*$^{nu}$ athymic nude mice were purchased from Charles River for TOV21G xenograft experiments. 5 x $10^5$ of TOV21G+EV or TOV21G+FANCF cells in PBS were mixed with equal volume of Matrigel (Corning) and injected subcutaneously to the flank of the mice. The mice were then randomly assigned to 4 treatment groups: (1) TOV21G+EV:PBS, (2) TOV21G+EV:NVB, (3) TOV21G+FANCF:PBS, and (4) TOV21G+FANCF:NVB. 4 days after tumor cell implantation, the mice were treated with NVB (100 mg/kg) or PBS twice a day via intraperitoneal injection (IP injection) for 4 weeks. Tumors were measured every 2 to 3 days using an electronic caliper, and tumor volumes were calculated by using the formula L×W×W/2.

**GEMM:** Pieces from breast tumors generated in K14-*Cre-Brca1*$^{f/f}$*Trp53*$^{f/f}$ female mice were transplanted into the mammary fat pad of FVB/129P2 recipient females that were at least 6 weeks old. FVB/129P2 recipients were generated by breeding FVB females (Jackson Laboratories) and 129P2 males (Envigo) and using the first-generation litters for experimentation. When tumors reached approximately 150 $mm^3$ in volume mice were randomized into treatment groups (vehicle and NVB). Treatments continued until tumors reached 20 mm in any direction, at which point mice were euthanized. NVB was prepared fresh from powder in PBS each time before IP injection and administered at 100 mg/kg twice a day for 35 days. Tumors were measured every 3 to 4 days using an electronic caliper, and tumor volumes were calculated by using the formula L×W×W/2.

**PDX studies:** Tumor ascites were collected from patients with suspected or established ovarian cancer at the Brigham and Women's Hospital or the Dana-Farber Cancer Institute (DFCI) under IRB-approved protocols conducted in accordance with the Declaration of Helsinki and the Belmont Report. Written informed consent was obtained from patients when required. PDX models were established, luciferized and propagated at the DFCI and have been described previously (*3*). Approximately 2-10 x $10^6$ luciferized PDX cells derived from mouse ascites were injected intraperitoneally into 8-week old female NSG mice. Tumor burden was measured by bioluminescence imaging (BLI) using a Xenogen IVIS-200 system (Xenogen). Mice were grouped according to their BLI signal from tumors so that each group would have similar initial average BLI. NVB was diluted in saline and administered via IP injection at 75 mg/kg twice daily for 4 weeks. Olaparib (ChemExpress) was formulated in PBS containing 10% DMSO and 10% (wt/vol) 2-hydroxypropyl- β-cyclodextrin (Sigma-Aldrich) and administered orally at 50 mg/kg daily for 4 weeks.

**Quantitative RT-PCR**

Total RNA was extracted using the RNeasy Mini kit (Qiagen), and cDNA was generated using the SuperScript™ III First-Strand Synthesis kit (Thermo Fisher). The PowerUp SYBR Green Master Mix (Thermo Fisher) reagent was used to run quantitative PCR on a QuantStudio 7 Flex Real-Time PCR System (Applied Biosystems). Two set of POLQ primers were used which gave same POLQ mRNA measurements. POLQ primer set 1 (forward: 5-TATCTGCTGGAACTTTTGCTGA-3; reverse: 5-

CTCACACCATTTCTTTGATGGA-3); POLQ primer set 2 (forward: 5-CTACAAGTGAAGGGAGATGAGG-3; reverse: 5-TCAGAGGGTTTCACCAATCC-3). Internal control primers are beta-Actin (forward: 5-CACCATTGGCAATGAGCGGTTC-3; reverse: 5-AGGTCTTTGCGGATGTCCACGT-3) or GAPDH (forward: 5-GTCTCCTCTGACTTCAACAGCG-3; reverse: 5-ACCACCCTGTTGCTGTAGCCAA-3).

**Immunohistochemistry (IHC)**

NSG mice bearing DF-59 and DF-83 tumors were dosed as in the efficacy study, with 75 mg/kg NVB twice a day and 50 mg/kg Olaparib daily. 4 hours after last dose (52 hours after first dose), tumor cells were isolated from the peritoneum of the mice, cleaned, washed with PBS and fixed in 10% normal buffered formalin for 10 min at RT. Next, the tumor cells were washed with PBS 3X and embedded in Histogel (Richard Alen Scientific). Histogel cores were processed using standard histology methods and embedded in paraffin. RAD51 staining was performed on paraffin sections (4 µm) using a previously standardized protocol (https://doi.org/10.1158/1538-7445.AM2017-2796). Images of stained slides were acquired on an Olympus BX41 microscope equipped with a digital camera at 40X magnification. The percentage of RAD51-foci positive cells fields was estimated.
